# Scaffold-Mediated Cellular and Acellular Delivery of Inducible BMP2-Lentivirus Promotes Atrophic Nonunion Fracture Healing without Heterotopic Ossification

**DOI:** 10.64898/2026.11.18.767681

**Authors:** Leyi Chen, Yu Seon Kim, Katherine R. Hixon, Nicole R. Gould, Andre F. Coello, Mariam Obaji, Tiffany Lee, Farshid Guilak, Matthew J. Silva

## Abstract

Nonunions occur in 5-10% of fracture patients, and current treatments are expensive with variable success. Controllable, localized delivery of BMP2 using porous, degradable scaffolds may be a promising alternative. We designed tube-shaped cryogel scaffolds (chitosan-gelatin, CG; chitosan-agarose-gelatin, CAG) to deliver a doxycycline-inducible BMP2-lentivirus vector (BMP2-LV). Results showed that CAG scaffolds have larger pores, higher toughness, and better cell infiltration than CG scaffolds. When scaffolds were implanted subcutaneously in mice and treated with continuous doxycycline, x-ray radiographs and µCT showed mineralization in scaffolds seeded with adipose-derived stem cells (BMP2-LV-ASC) but not acellular scaffolds (BMP2-LV) 4-weeks after implantation. Scaffolds that were implanted at the site of an established femoral nonunion 4-weeks after fracture and treated with continuous doxycycline showed improved healing with CG/CAG BMP2-LV and BMP2-LV-ASC compared to scaffold-only controls. At 2-weeks post implantation, µCT and histology showed that BMP2-expressing scaffolds promoted increased callus mineralization relative to scaffold-only. At 8-weeks post implantation, µCT and histology showed near-complete bone bridging in BMP2-treated groups compared to still visible fracture lines in the scaffold-only groups. However, CG/CAG BMP2-LV-ASC groups also showed heterotopic ossification. Finally, we demonstrated that heterotopic ossification was eliminated at 8-weeks post implantation in CG BMP2-LV-ASC groups by restricting doxycycline treatment to the first 3 days after implantation. We conclude that lentiviral BMP2 delivery using cellularized and acellular cryogel scaffolds enhanced fracture healing in a nonunion model, and that a 3-day period of BMP2 induction using an inducible system was sufficient to promote healing without the complication of heterotopic ossification.

## Introduction

Bone is one of the few tissues in the human body with the inherent capacity to regenerate without forming scar tissues and to fully restore its original function. However, under circumstances of insufficient mechanical stability or impaired biological activity, a fracture may not be able to heal, resulting in a fracture nonunion that requires further intervention[1]. Clinically, the overall risk of nonunion is estimated at approximately 5% among all fractures, with treatment costs reaching up to approximately $60,000/patient/year[2,3]. Atrophic nonunion is a subtype of nonunions characterized by insufficient osteochondral potential[1]. The underlying biological mechanisms driving atrophic nonunion remain poorly understood, which contributes to the challenges in its clinical treatment. Currently, an autologous bone graft with internal fixation is considered to be the gold standard for treating atrophic nonunion[4]. However, this approach is limited by finite graft volume, donor-site morbidity including chronic pain, risk of infection at both surgical sites, and variable osteogenic potency in elderly or comorbid patients[5,6].

Atrophic nonunion models have been established in the mouse and rat via stripping the periosteum along with osteotomy at the tibial or femoral midshaft[7–10]. These models are relevant to conditions of extensive soft tissue disruption, but are not relevant to the cases where nonunion arises without extensive tissue damage. To model the less traumatic cases where atrophic nonunions are due to “failure of biology”, we treated 3.6Col1a1-TK (Col1-TK) transgenic mice with ganciclovir (GCV) to selectively ablate proliferating osteoblast lineage cells[11]. Blocking osteoblastic cell proliferation for 2 weeks after fracture in Col1-TK mice resulted in the failure of mineralized callus formation leading to atrophic nonunion that persisted up to 12 weeks post-fracture [12].

Biomaterial scaffolds are among the most widely studied alternatives to autograft due to their highly adjustable material and mechanical properties [4]. Cryogels are a subset of hydrogel scaffolds with porous microstructure and enhanced mechanical properties as a result of a freeze-thaw fabrication process [13,14]. In combination with appropriate cell types and/or osteoinductive factors, such as BMP2 [15], tricalcium phosphate (TCP) [16], or vascular endothelial growth factor (VEGF)[17], cryogel scaffolds have considered to be promising candidates for bone graft substitutes. The clinically available synthetic bone graft, INFUSE^®^, is a combination of recombinant human BMP2 (rhBMP2) protein and absorbable collagen sponge and has been FDA approved for use in spinal fusion surgery since 2007[18,19]. However, the side effects of uncontrollable supraphysiological doses of rhBMP2 protein, such as extensive swelling, bone overgrowth, and cancer development remain concerns[19,20]. In addition, protein-based therapeutics have relatively short half-life and high clearance rate *in vivo*, limiting their suitability for sustained release and necessitating repeated dosing for long-term fracture healing[19,21].

Gene delivery systems offer a promising alternative for achieving sustained release of proteins *in vivo*[21]. Through the use of a delivery vector, either viral or non-viral, specific genetic material can be delivered into the target cells, leading to transgene expression and secretion of the desired biomolecules. Lentivirus (LV), compared to the other viral vectors, such as adenovirus (Ad) and adeno-associated virus (AAV), provides the most long-term and stable gene expression with low immunogenic activity[21,22]. Previous studies have demonstrated the ability of LV-based methods to mediate stem cell differentiation and cartilaginous or mineralized extracellular matrix (ECM) formation on biomaterial scaffolds[23–25]. Recently, we developed a chitosan-gelatin (CG) cryogel scaffold for the delivery of doxycycline-inducible BMP2-LV-transduced adipose-derived stem cells (ASC), and preliminary studies showed promotion of bone formation in the Col1-TK atrophic nonunion model[15]. However, CG scaffolds are fragile making surgical implantation difficult, and the pore size of the chitosan-gelatin (CG) cryogels in the prior study was limited to ∼30µm, which is smaller than optimal for supporting angiogenesis and cell infiltration[15]. In addition, excessive heterotopic ossification was observed in the BMP2-LV-ASC group, highlighting the need for further optimization of scaffold design and delivery strategies [15].

Agarose, a natural polysaccharide, has been widely used as the matrix material for bone tissue engineering[26,27]. In this study, agarose was incorporated with chitosan and gelatin to form chitosan-agarose-gelatin (CAG) cryogel scaffolds. We hypothesize that the addition of agarose results in larger pore size and higher porosity, thereby promoting cell infiltration, while also enhancing mechanical properties. To achieve the controllable and localized treatment of atrophic nonunion, we evaluated the osteogenesis potentials of both CG and CAG scaffolds as delivery matrices for BMP2-LV delivery through both cellular (BMP2-LV-ASC) and acellular (BMP2-LV) delivery strategies in the Col1-TK murine nonunion model. The BMP2-LV was constructed with a doxycycline-inducible promoter[28], allowing the magnitude and duration of BMP2 production to be tuned by adjusting the doxycycline dose administered. We hypothesize that localized, scaffold-mediated gene-delivery of osteogenic transgene BMP2 induces osteogenesis at the nonunion fracture site, while the acellular (BMP2-LV) delivery and dose-adjustable transgene expression reduces the risk of excessive heterotopic ossification.

## Methods

### Scaffold Fabrication

Cryogel scaffolds were fabricated based on a previous protocol[15]. Briefly, for chitosan-gelatin scaffolds (CG), 80mg chitosan (MP Biomedicals) were dissolved in 1% acetic acid solution on a mechanical spinner (MIDSCI). Subsequently, 320mg gelatin (from cold water fish skin; Sigma) was added to the chitosan solution and mixed until homogeneous. Chitosan-agarose-gelatin scaffold (CAG) fabrication was based on a published protocol[29] except that the composition used here is 100mg chitosan, 300mg agarose and 100mg gelatin. For all scaffolds, glutaraldehyde (CG: 1%; CAG: 0.2%) was prepared separately in a scintillation vial as the crosslinker. Solutions were then precooled under 4°C for at least 1h. The polymer and the crosslinker were mixed by decanting and then injected into customized 3D printed molds (Supplemental Fig. S1; Prusa Research) using 0.3cc syringes (BD) with 18G needles (BD). The scaffolds were designed to be tube shaped to facilitate placement around the mouse femur. The polymer-filled molds were frozen to -20°C as follows. For the CG scaffolds, the molds were frozen in a -20°C methanol bath for at least 18h. For CAG scaffolds, to achieve different pore sizes the molds were frozen by three different methods to vary the rate of cooling: in a methanol bath (rapid freeze; small-pore, SP), in air (slower freeze; medium-pore, MP), and in a sponge thermal isolator in air (slowest freeze; large-pore, LP) for at least 18h. These are referred to as CAG-SP, CAG-MP and CAG-LP, respectively. Molds were then thawed in water at room temperature (RT), scaffolds were removed and lyophilized and stored at RT until use. Before adipose-derived stem cell (ASC) and lentivirus (LV) seeding, scaffolds were sterilized in 1% peracetic acid (PAA; Sigma) for 1.5h and then rinsed with sterile 1x phosphate-buffered saline (PBS; ThermoFisher) three times for 10 min and lyophilized overnight. All the scaffolds for implantation were trimmed to 5mm in length.

### Tensile Testing

CG and CAG-SP scaffolds were trimmed into a rectangular shape (2mm x 0.5mm x 20mm) and evaluated by mechanical testing. Lyophilized scaffolds (n=6/group) were rehydrated in water for 1 min right before loading to failure using a tensile testing machine (ElectroForce 3200; TA Instrument). The scaffold was fixed by tensile grips (Fig. 1A). A 1000gram (9.8N) load cell was used to record force during testing. The initial gauge length (L₀), defined as the free specimen length between the inner faces of the upper and lower grips, was measured for each sample prior to testing (6.5±1mm). Strain (ε) was calculated as *ε* = *ΔL*/*L*_0_ while the stress (σ) was determined by *σ* = *F*/*A*. Samples were stretched under displacement control at 0.05mm/s. A customized MATLAB code was used to generate the stress-strain curves from which we determined yield stress, ultimate stress, Young’s Modulus (E) and Post Yield Toughness based on the following equations:

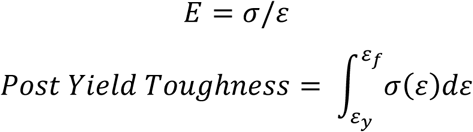

**Figure 1.**
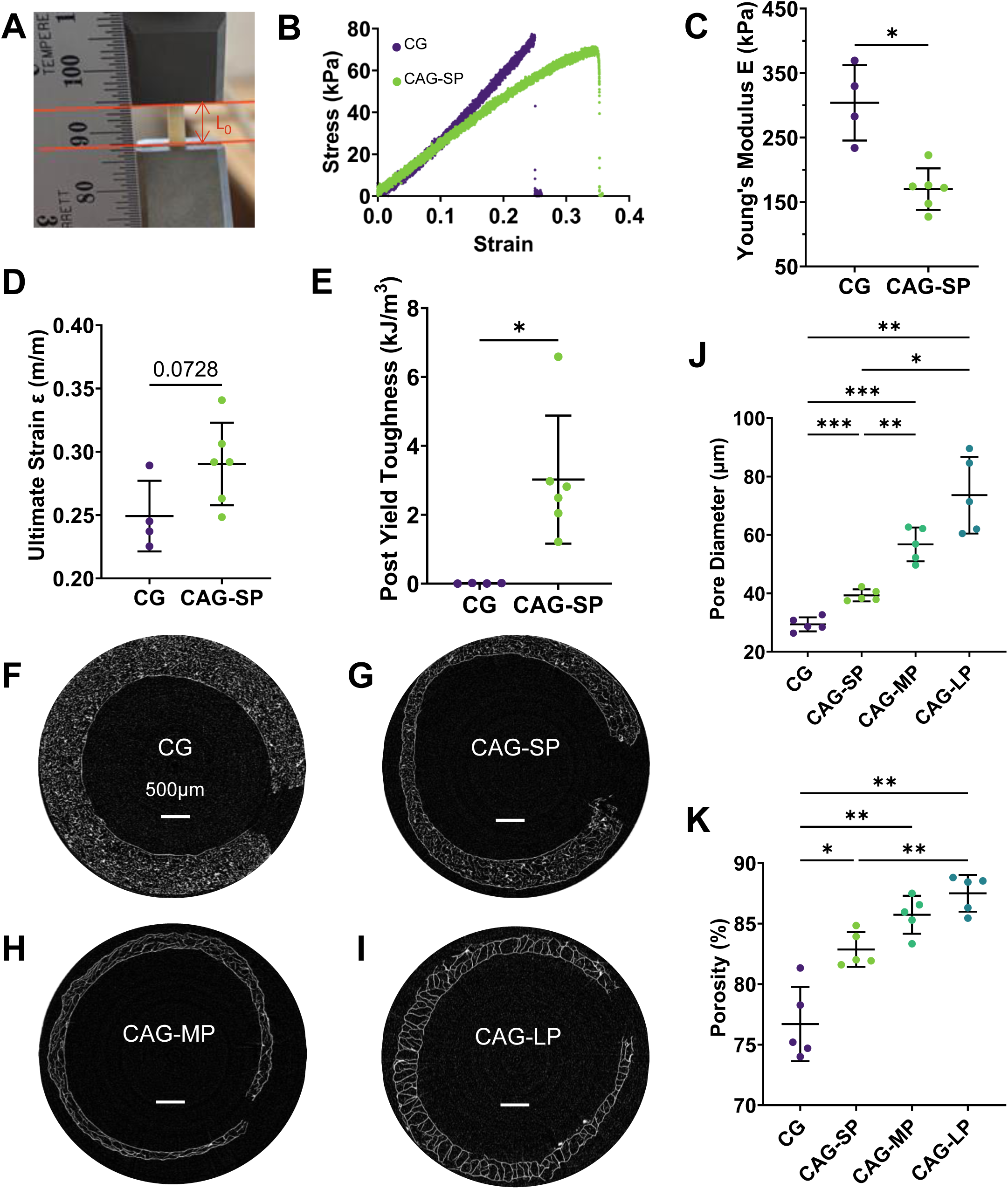
Mechanical and structural characterization of chitosan-gelatin (CG) and chitosan-agarose-gelatin (CAG) cryogel scaffolds. **(A)** Uniaxial tensile testing setup. Red lines indicate the inner grip faces defining the initial gauge length (L₀). (B) Representative stress-strain curves for CG and CAG-SP scaffolds. (C) Young’s modulus, (D) ultimate strain, and **(E)** post-yield toughness of CG (n=4) and CAG-SP (n=6) scaffolds. **(F-I)** Representative µCT cross-sections of **(F)** CG, **(G)** CAG-SP, **(H)** CAG-MP, and **(I)** CAG-LP scaffolds. Scale bars, 500µm. **(J)** Pore diameter and **(K)** porosity quantified from µCT (n=5/group). SP, MP, and LP denote small, medium, and large pore formulations. Data are mean ± SD. Statistical comparisons were performed by unpaired t-test (C-E) and one-way ANOVA with Tukey’s post-hoc test (J, K). *p < 0.05, **p < 0.01, ***p < 0.001.

Where ε_y_ is the strain at yield, ε_f_ is the strain at fracture. The yield point was automatically identified based on the criterion where the stress–strain curve began to deviate from linear elastic behavior. The scaffolds that were fractured at the grips were excluded from analysis.

### Scaffold Micro-Computed Tomography

Following lyophilization, the scaffolds (n=5/group) were scanned in air by μCT (μCT 50; Scanco) at 45kV, 88μA, 3.4μm voxel size. 540 slices were scanned for each scaffold and a single threshold (50 per milles) was used to determine the microstructure of the scaffolds based on manufacturer installed trabecular morphology analysis. Specifically, the average pore diameter (μm) was determined by trabecular bone separation (Tb.Sp) while the porosity was determined by bone volume over total volume (BV/TV) [30].

### Lentivirus Production

Second-generation lentivirus expressing doxycycline (dox)-inducible BMP2 and constitutive GFP expression vectors were produced using an established protocol[15]. Briefly, 293T/17 cells (ATCC CRL-11268) were seeded onto 15-cm petri dishes at 3 million cells/ml density in growth medium, composed of 10% fetal bovine serum (FBS, Gibco) and 1% penicillin/streptomycin (Gibco) in high-glucose Dulbecco’s Modified Eagle Medium (DMEM, Gibco), and allowed to grow to 80-90% confluency. Cells were transfected with second generation lentiviral packaging and envelope plasmids (psPAX2, Addgene #12260; pMD2.G, Addgene #12259) and expression plasmid (dox-inducible BMP2 or constitutive GFP vector) via calcium phosphate precipitation method[31]. Media was changed 16 h post-transfection, and supernatant was collected 24 and 48h post-media change. Collected media was concentrated ∼70-fold using 100kDa molecular weight-cutoff centrifuge filters. Lentivirus concentrate was aliquoted in 2-ml cryovials and stored in -80 ℃ until further use.

### Adipose-Derived Stem Cell Isolation

To prepare cellular scaffolds (BMP2-LV-ASC), adipose-derived stem cells were isolated from 12-week-old C57BL6/J mice (Jackson Lab) based on a previously published protocol [15]. Specifically, subcutaneous adipose tissue was dissected from mice immediately after euthanasia (CO_2_ asphyxiation) under sterile conditions. The dissected tissue was minced and incubated in an enzymatic digestion solution containing 0.1% Collagenase Type I (Gibco) and 1% penicillin/streptomycin in DMEM for 1h at 37°C with mechanical inversion every 10min. Following digestion, the tissue was transferred to a culture dish and dissociated using forceps, followed by a 30-minute incubation in 37°C incubator. At this time, the tissue was further dissociated by pipetting, and then filtered through a 70μm cell strainer and centrifuged at 1300rpm for 10min at 4°C. The supernatant was resuspended in the growth medium and immediately plated into a T150 flask. All the adipose-derived stem cells were seeded on the scaffolds before passage five for *in vivo* implantation.

### Scaffolds-Mediated Lentiviral Transduction

Scaffolds were incubated overnight with 0.002% poly-L-lysine (PLL, Sigma), rinsed with 1x PBS, and then incubated with lentivirus (∼ 5 x 10^5^ viral particles/scaffold) for 1.5h at 37°C in a low-attachment 24-well plate (Corning). The scaffolds were then rinsed again with 1x PBS. For cellularized groups, scaffolds were seeded with 500,000 cells for *in vitro* experiments and 1,000,000 cells for *in vivo* experiments and incubated for 1h under 37°C before adding the growth medium. The growth medium was changed every 2 days for 5 days to facilitate cell proliferation before switching to osteogenic medium containing 50ug/ml L-ascorbic acid 2-phosphate (Sigma), 10nM dexamethasone (Sigma) and 10mM β-glycerophosphate (Sigma) in growth medium. The timeline of scaffold preparation is shown in Supplemental Figure S2.

### Scaffolds Preparation for Implantation

Four scaffold types and three delivery pathways were prepared for *in vivo* implantation: scaffold only (CG, CAG-SP, CAG-MP and CAG-LP), scaffold + BMP2-lentivirus (BMP2-LV), and scaffold + ASCs transduced to express BMP2 (BMP2-LV-ASC). In preparation for *in vivo* implantation, BMP2-LV-ASC scaffolds underwent *Scaffold-mediated Lentiviral Transduction* as described above and incubated in osteogenic medium for 10 days. Scaffold only and BMP2-LV groups were coated with PLL for 24h before the implantation. The BMP2-LV scaffolds were implanted within 2h once the BMP2-lentivirus was immobilized on the scaffolds.

### In vitro BMP2 Production

To evaluate BMP2 production *in vitro*, CG/CAG-SP scaffolds ± BMP2-lentivirus transduced ST2 stromal cells were cultured in osteogenic medium with or without administration of 1μg/ml doxycycline (dox, Sigma). Half medium changes were performed every other day for 10 days, and the media samples were collected on days 0, 2, 6 and 10. The concentration of BMP2 in the media (n=3/group/timepoint) was determined by ELISA (R&D DY355).

### Subcutaneous Implantation Model

Experimental procedures were approved by the Institutional Animal Care and Use Committee (IACUC) at Washington University in St. Louis. Scaffolds were implanted subcutaneously on the dorsum of 12-week-old C57BL/6J mice (Jackson Lab). Specifically, each mouse had four scaffold pieces implanted, which were trimmed into rectangular shape (2mm x 0.5mm x 5mm) (Fig. 2A). All mice were treated with Buprenorphine SR (1mg/kg, s.c.) within 2h before surgery. After the implantation, each mouse was single housed to protect the wounds. Weekly radiography was performed to track any mineralization in the region of the implanted scaffolds.

**Figure 2.**
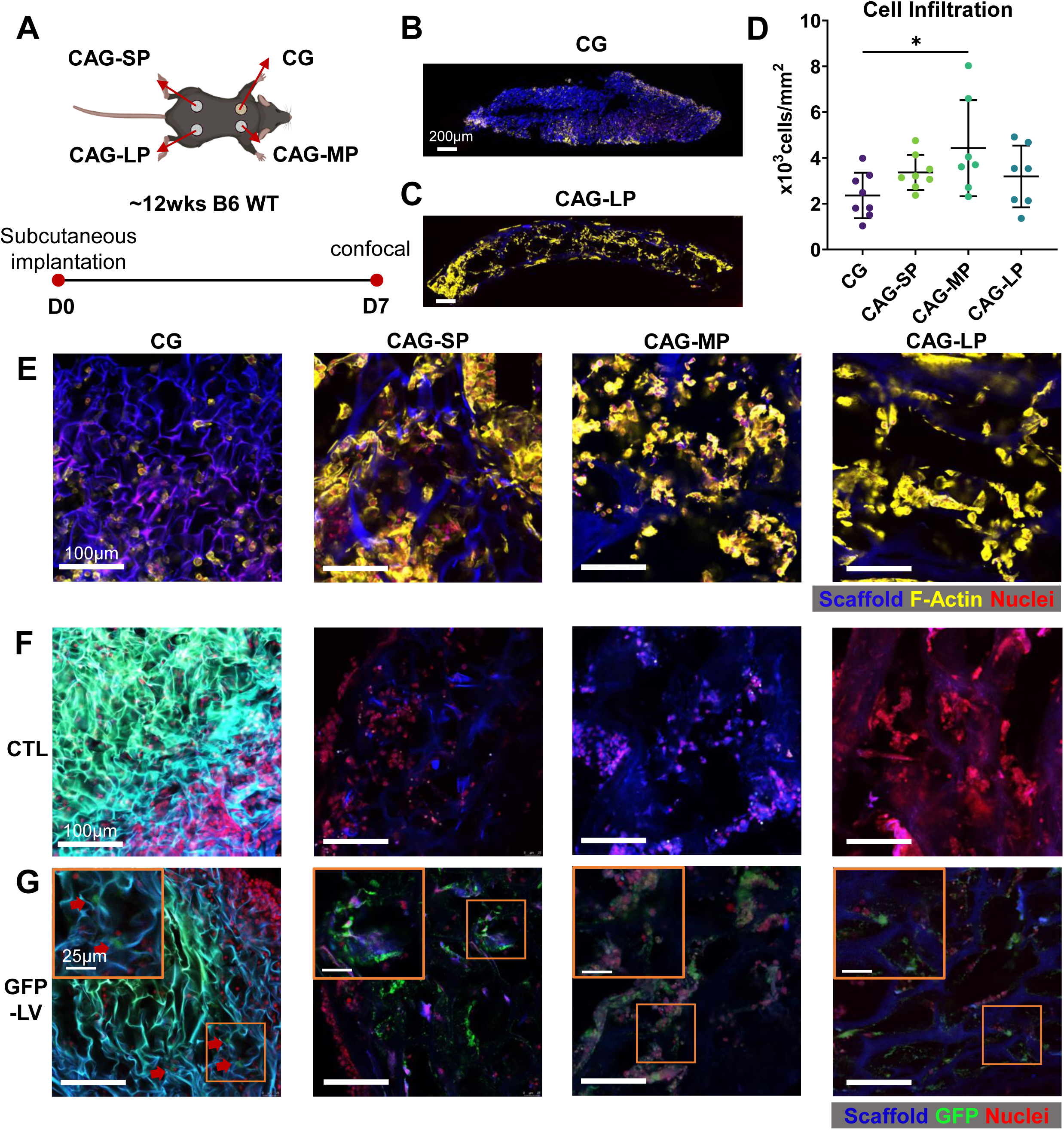
Confocal microscopy illustrated cell infiltration and lentiviral (LV) transduction in CG and CAG scaffolds in a subcutaneous implantation model. **(A)** Experimental schematic. CG, CAG-SP, CAG-MP, and CAG-LP scaffolds were implanted subcutaneously in ∼12-week-old C57BL/6 mice and harvested at day 7 for confocal imaging. **(B, C)** Representative cross-sections of (B) CG and (C) CAG-LP scaffolds without lentivirus. Scale bar, 200µm. **(D)** Quantification of cell infiltration (n=6-8/group). (E) Representative confocal images of cell infiltration in each scaffold formulation. Scaffold autofluorescence (blue), F-actin (yellow), and nuclei (red). Scale bars, 100 µm. **(F, G)** Representative confocal images of scaffolds implanted without virus **(F, CTL)** and with GFP-lentivirus **(G, GFP-LV)**. Scaffold autofluorescence (blue), GFP (green), and nuclei (red). Insets show higher-magnification views of the boxed regions; inset scale bar, 25µm. Scale bars, 100µm. Data are mean ± SD. Statistical comparison by one-way ANOVA with Tukey’s post-hoc test. *p < 0.05.

### Confocal Microscopy

Confocal microcopy (Leica DMi8) was performed to detect *in vivo* cell infiltration (n=6-8/group) and LV transduction (n=3/group) in the subcutaneous model. To evaluate transduction, scaffolds were coated with PLL overnight and loaded with or without GFP-lentivirus (∼5 x 10^5^ viral particles/scaffold) within 2 h before subcutaneous implantation. GFP was used in place of BMP2 because it is retained intracellularly and directly visualized, allowing visualization of transduced cells within the scaffold. Each mouse was implanted with one each CG, CAG-SP, CAG-MP and CAG-LP. The scaffolds were harvested 1 week after implantation, fixed with 4% paraformaldehyde (PFA), and cryoprotected with 30% sucrose in PBS overnight at 4°C before cryosection. Next, the scaffold sections were stained with NucRed^TM^ Live 647 ReadyProbes^TM^ Reagent for nuclei (Invitrogen^TM^) and Alexa Fluor™ 555 Phalloidin for F-actin (1:1000, Invitrogen^TM^). A tile scan was performed to scan the whole scaffold piece for each sample. Three regions (150µm x 150µm) were randomly selected for each sample for cell counting using QuPath^®^.

### qPCR

Four scaffold types (CG, CAG-SP, -MP and -LP) and three delivery methods (BMP2-LV-ASC, BMP2-LV, and scaffold only) were used to characterize *Bmp2* mRNA expression via qPCR. The scaffolds were dissected 2 weeks after implantation and snap frozen by liquid nitrogen. After that, the scaffolds were pulverized, and mRNAs were isolated by TRIzol (Invitrogen). Primer Time assay (Integrated DNA Technologies) Mm.PT.58.10419414 was used to detect *Bmp2* expression detect in the scaffold. The assay amplifies the region spanning exons 2 and 3. Mm.PT.39a.22214839 was used to amplify exons 4-5 to detect *Tbp* expression as a reference gene.

### Atrophic Nonunion Fracture Model

Experimental procedures were approved by the Institutional Animal Care and Use Committee (IACUC) at Washington University in St. Louis. Atrophic nonunion fractures were established in 12-week-old transgenic 3.6Col1a1-TK (Col1-TK) mice by targeting proliferating osteoblasts[12]. Specifically, the herpes simplex virus thymidine kinase gene (HSV-TK, or “TK”) was driven by the 3.6Col1a1 promoter. TK transforms the guanosine analogue ganciclovir (GCV) into ganciclovir-triphosphate. This toxic version of a nucleotide will then incorporate into DNA of dividing cells and lead to cell apoptosis[32]. All Col1-TK^+^ mice used in the experiments are heterozygous because male Col1-TK^+^ mice are sterile. A previously established protocol was used to create a full femur fracture in the right femur of 12-week-old mice[33]. All mice were treated with Buprenorphine SR (1mg/kg, s.c.) 2h before surgery. The fracture was performed using a customized 3-point bending platform by applying a transverse force at the right femoral mid-diaphysis (Dynamite 8841; Instron). A 24-gauge metal pin (MicroGroup) was then used to stabilize the fracture, and 4-0 nylon sutures (Ethicon) were used to close the incision. X-ray radiography (Faxitron Ultrafocus 100) was performed immediately after the fracture to confirm proper fracture site and stabilization. To establish atrophic nonunion, GCV (8mg/kg, i.p) was dosed once/day for 2 weeks starting from the fracture day. Weekly radiography was performed to confirm minimal callus formation and healing response until 4 weeks after fracture surgery, consistent with a nonunion. Two Col1-TK^-^ (WT) mice were included in each surgery batch to check for normal healing in WT mice, confirming that GCV only inhibits callus formation in Col1-TK^+^ mice.

### Scaffold Implantation

Two scaffold types (CG and CAG-MP) and three delivery methods (BMP2-LV-ASC, BMP2-LV and scaffold only) were used for scaffold implantation in atrophic nonunion model. Specifically, the scaffold was implanted 4 weeks after fracture once the atrophic nonunion was confirmed radiographically in Col1-TK^+^ mice. All mice were administered with Buprenorphine SR (1mg/kg, s.c.) within 2h before the surgery. To expose the fractured femur, an incision was made along the femur, and the muscle was gently separated from the nonunion site. The scaffold (n=5-10/group/timepoint) was placed under the femur, wrapped around the fracture site. Then, the scaffold was held in position and the muscle was closed with 5-0 vinyl sutures (Ethicon). After that, the incision was closed using running subcuticular 6-0 PDS-II sutures (Ethicon). To activate transgene expression, mice started to receive doxycycline chow (Inotiv, 625mg/kg) 2 weeks before fracture surgery (D0) and changed weekly. In continuously induced groups, doxycycline was maintained until sacrifice. In the shortened-induction group, doxycycline was withdrawn 3 days after scaffold implantation at week 4. Because the BMP2-LV construct is not present in the animal until implantation, this regimen restricts transgene expression to a three-day window. Weekly radiographs were performed to track callus formation after the surgery. All mice were euthanized 2 or 8 weeks after implantation based on previous studies[15,34]. Both the left and right femurs were dissected for µCT and histological staining.

### Radiographic Evaluation

Weekly radiographs were taken after the fracture until euthanasia to track the fracture healing (n=5-10/group/timepoint). All radiographs were blindly scored by a single reader using the modified mRUST scoring system for the degree of healing. Both cortical sides were scored individually from score 1 (only the fracture line) to score 4 (bridging with no visible fracture line) with a maximum score of 8 in total[35]. Any mice that lost the pin during the healing process were excluded from further analysis.

### Micro CT

All mice were euthanized at 2 or 8 weeks after scaffold implantation. Both the left and right femurs were dissected and fixed with 10% neutral buffered formalin (NBF) for µCT scanning (µCT 50, Scanco; 6μm voxel size, 70kV, 57μA). The pin was removed after fixation. To evaluate the properties of the callus of each right fractured femurs, the outer boundary of the callus was contoured semi-automatically using a threshold of 450mgHA/cm^3^ to include the total bone area. Then, the boundary of the cortical bone was contoured using the same threshold to exclude the original cortical bone and the marrow. The contour was morphed every 10-20 slices to a maximum of 1050 slices (6.3mm length) centering at the middle of the callus. The callus TV, BV, BV/TV, BMD and TMD were measured automatically by a manufacturer installed trabecular morphology analysis using a threshold of 450mgHA/cm^3^. Any bone formation that is completely outside of the normal skeleton or not incorporated into the original tissue was considered as heterotopic bone formation. The boundary of the heterotopic bone formation was semi-automatically contoured using a threshold of 300mgHA/cm^3^ and morphed every 10-20 slices. A manufacturer installed trabecular morphology analysis program was used to measure the heterotopic BV using a threshold of 300mgHA/cm^3^.

### Histological Staining and Analysis

The samples were decalcified with 14% EDTA (pH=7) for 10 days (n=5-10/group/timepoint) after µCT. Then, the samples were embedded in paraffin and sectioned longitudinally at 5µm thickness. The slides were stained with picrosirius red/alcian blue (PSRAB) to visualize the tissue composition of the callus, including mineralized woven bone, cartilage, and fibrous tissue. The slides were scanned (nanozoomer, Hamamatsu) and analyzed by a blinded observer (NDP.view, Hamamatsu) to determine the %area of each tissue.

### Statistics

Sample size was determined by a power analysis with α = 0.05 and β= 0.20 (https://www.stat.ubc.ca/~rollin/stats/ssize/n2.html) based on our prior results and expected effect sizes. An unpaired t-test was used for scaffold mechanical properties analysis. One-way ANOVA with the Tukey *post hoc* was used for scaffold microstructure, cell infiltration, and subcutaneous mineralization. Kruskal-Wallis test with Dunn’s *post hoc* and Chi-square test for trend were used to compare mRUST radiographic scoring at 2- and 8-weeks post-scaffold implantation. Two-way ANOVA with the Tukey *post hoc* was used for µCT and PCR multiple comparison. P-value < 0.05 was considered as a significant difference. The data analysis was performed via Prism (GraphPad ver.10).

## Results

### CAG Scaffolds Exhibited Enhanced Toughness and Porosity Compared to CG Scaffolds

Lyophilized CAG and CG scaffolds were trimmed into rectangular shapes (2mm x 0.5mm x 20mm) and rehydrated for tensile testing (Fig. 1A). The stress and strain curves revealed distinct mechanical behaviors between the two scaffold types (Fig 1B). Specifically, CG scaffolds exhibited a 79% higher Young’s modulus (p < 0.05; Fig. 1C) and a 48% higher ultimate stress (p < 0.05; Supplemental Fig. S3A). Conversely, CAG scaffolds demonstrated greater ductility, with marginally higher ultimate strain (*p* = 0.073; Fig. 1D) and markedly increased post-yield toughness (3.02±0.4kJ/m³ vs. 0.01±0.01kJ/m³, *p* < 0.05; Fig. 1E). Notably, CG scaffolds did not exhibit a yield point before failure (Fig. 1B, Supplemental Fig. S3B, C), indicating a more brittle mechanical behavior compared to CAG scaffolds. The total absorbed energy was comparable between groups (Supplemental Fig. 3D).

Microcomputed tomography (μCT) revealed distinct microarchitectures between CG and CAG scaffolds (Fig. 1F-I). Specifically, CG scaffolds showed significantly lower porosity (76.7±3.1%) and smaller pore size (26.4±2.4μm) than CAG scaffolds (Fig. 1J, K). Furthermore, within the CAG groups, both porosity and pore size increased progressively with decreasing freezing rate, with the CAG large-pore (LP) exhibiting the highest porosity (87.5±1.5%) and largest pore size (73.6±13.1μm; Fig. 1J, K).

### CAG Scaffolds Demonstrated Enhanced Cell Infiltration and Lentiviral (LV) Transduction Efficiency Compared to CG Scaffolds

Lyophilized CAG and CG scaffolds were implanted subcutaneously for *in vivo* evaluation (Fig. 2A). 7 days after subcutaneous implantation, confocal imaging revealed markedly greater cellular infiltration in CAG scaffolds than in CG scaffolds (Fig. 2B, C). Quantitative analysis confirmed the highest cell density in CAG MP scaffolds (4.43×10³±2.10×10³cells/mm²), which significantly exceeding that in CG scaffolds (2.36×10³±0.99×10³ cells/mm²; *p* < 0.05; Fig. 2D). High-magnification imaging further validated deeper cell penetration within CAG scaffolds, while cells in CG scaffolds were primarily localized to the periphery (Fig. 2B, E).

To confirm LV transduction, scaffolds were treated with GFP-expressing lentivirus (GFP-LV) 1-2 h prior to implantation or left untreated (CTL). Confocal imaging revealed GFP⁺ cells distributed throughout both CG and CAG scaffolds, confirming effective *in vivo* lentivirus transduction of cells that have migrated into the scaffold (Fig. 2F, G).

### Cellular Delivery Pathway (BMP2-LV-ASC) Exhibited Extensive Mineralization in Subcutaneous Model

First, to validate the doxycycline-inducible BMP2 expression from BMP2-LV, ST2 bone marrow stromal cells were transduced with BMP2-LV or left untransduced and seeded onto CG or CAG small-pore (SP) scaffolds. Scaffolds were cultured *in vitro* with or without doxycycline and supernatants were collected at D0, D2, D6 and D10 for ELISA. Robust BMP2 production was detected only in doxycycline-treated BMP2-LV groups, whereas negligible BMP2 production was observed in all other conditions (Fig. 3A). BMP2 secretion remained stable over 10 days and was higher in CAG-SP scaffolds than CG scaffolds under doxycycline induction. (Fig. 3A)

**Figure 3.**
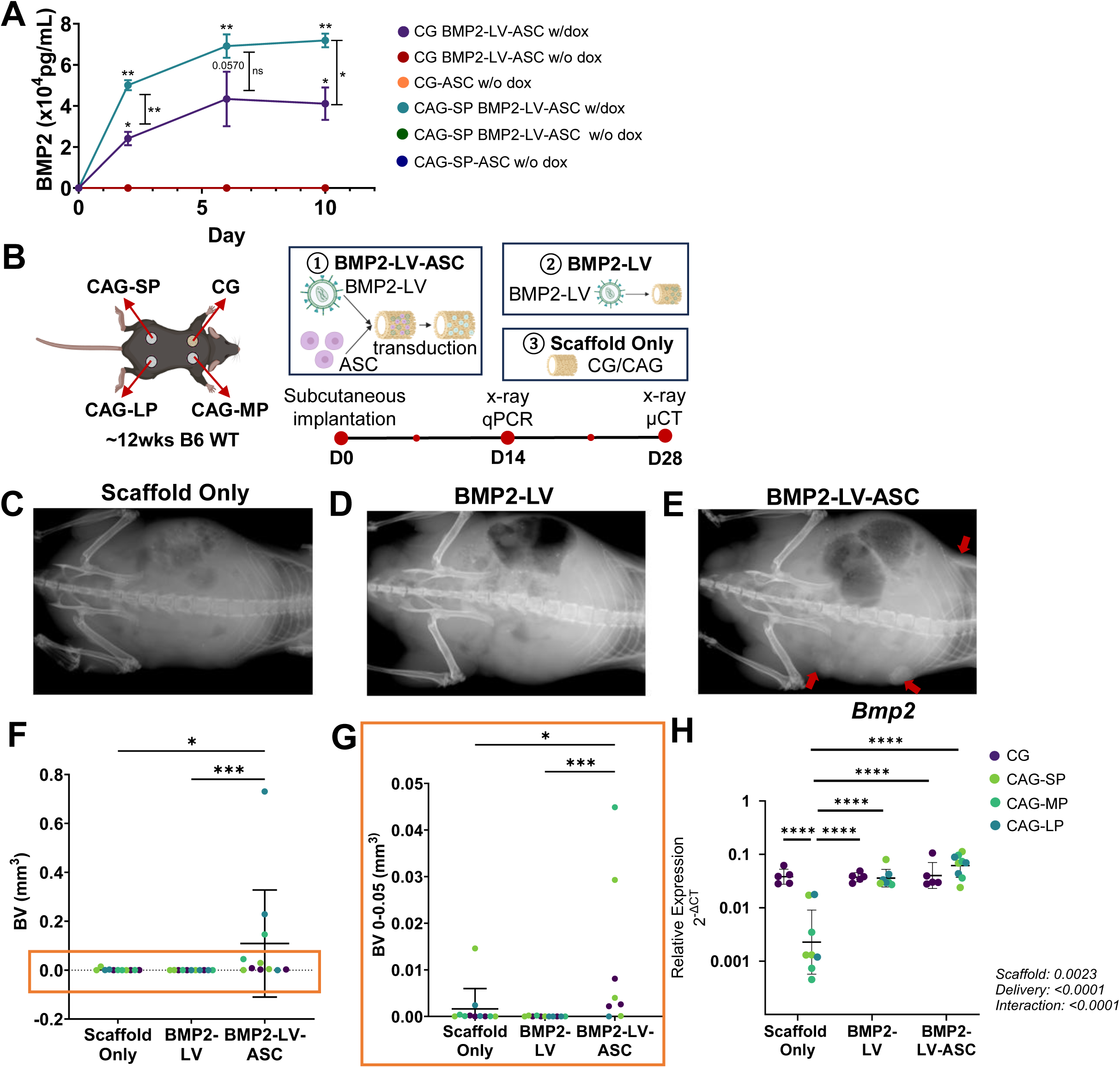
BMP2-expressing cryogel scaffolds exhibit transgene expression and bone formation in a subcutaneous model. **(A)** BMP2 concentration in culture medium measured by ELISA at days 0, 2, 6, and 10. Analysis by repeated-measurement two-way ANOVA with Geisser-Greenhouse correction. Asterisk markers adjacent to each curve indicate comparisons versus day 0 within that group (Dunnett’s test); vertical brackets indicate CG versus CAG-SP at the indicated timepoint (Šídák test). All groups have negligible value at baseline (day 0). *p < 0.05, **p < 0.01; ns, not significant. **(B)** Experimental schematic. CG, CAG-SP, CAG-MP, and CAG-LP scaffolds were loaded with BMP2-LV-transduced ASCs (BMP2-LV-ASC), BMP2-LV alone, or left unmodified (scaffold only) and implanted subcutaneously in ∼12-week-old C57BL/6 mice. (C-E) Representative radiographs at day 28 of **(C)** scaffold-only, **(D)** BMP2-LV, and **(E)** BMP2-LV-ASC implants. Red arrowheads indicate scaffold mineralization. **(F)** bone volume (BV) quantified by µCT at day 28. **(G)** Magnified view of the boxed region in (F). **(H)** Bmp2 expression in CG and CAG scaffolds under scaffold-only, BMP2-LV, and BMP2-LV-ASC conditions. Relative expression calculated as 2^-ΔCT with Tbp as the reference gene, displayed on a log₂ axis. Symbols represent individual samples (n=5-7/group); bars indicate geometric mean ± geometric SD. Two-way ANOVA was performed on ΔCT values, with Tukey post-hoc comparisons. ****p < 0.0001.

The osteoinductive potential of BMP2-expressing scaffolds was then evaluated *in vivo* using a subcutaneous implantation model (Fig. 3B). Four scaffold types (CG, CAG-SP, -MP and -LP) and three delivery strategies were assessed, including cellular scaffolds with BMP2-LV-transduced adipose-derived stem cells (BMP2-LV-ASC), acellular scaffolds with BMP2-LV (BMP2-LV) and scaffold-only controls. After 28 days of subcutaneous implantation, X-ray imaging revealed visible mineralization in the BMP2-LV-ASC scaffolds, whereas BMP2-LV and scaffold-only groups exhibited minimal or no mineral deposition (Fig. 3C–E). μCT quantification confirmed significantly greater bone volume (BV) in the BMP2-LV-ASC groups compared with negligible bone volume in the acelluar BMP2-LV group and scaffold-only group (*p* < 0.0001 and *p* < 0.05, respectively; Fig. 3F, G). Interestingly, qPCR results showed significant interaction between scaffold composition and delivery strategy (p < 0.0001; Fig. 3H). In the absence of virus vector, CG scaffolds expressed substantially higher endogenous *Bmp2* than CAG (p < 0.0001), whereas expression was comparable between scaffold types following delivery of acellular BMP2-LV or cellular BMP2-LV-ASC (Fig. 3H). Together, these results demonstrate that cellular BMP2-LV-ASC delivery enables sustained BMP2 expression and promotes scaffold mineralization in the subcutaneous model. Since mineralization and gene expression did not differ significantly among CAG-SP, -MP, and -LP scaffolds, subsequent experiments focused on CG and CAG-MP scaffolds.

### BMP2-expressing Scaffolds Promoted Radiographic Healing in Nonunion Fractures

Following ganciclovir-induced osteoblast ablation and doxycycline administration, WT and Col1-TK mice received CG or CAG-MP scaffolds loaded with BMP2-LV-ASC, BMP2-LV, or left unmodified, and healing was assessed by blinded weekly mRUST scoring for 6 or 12 weeks post fracture (2 or 8 weeks post scaffold implantation; Fig. 4A, B). As expected, WT mice achieved complete cortical bridging as early as 2 weeks after fracture (Supplemental Fig. 4). In contrast, fewer than 50% of Col1-TK mice implanted with scaffold only (CG or CAG-MP) reached an mRUST score of 6 (two cortices bridged) at 8 weeks after implantation (week 12 post-fracture), confirming the impaired fracture healing in Col1-TK mice (Fig. 4C, D). On the other hand, 100% of fractures in BMP2-LV and BMP2-LV-ASC groups achieved a score ≥ 6 in at the same timepoint (Fig. 4C, D). Both CG and CAG-MP BMP2-expressing scaffolds exhibited significantly higher mRUST scores than scaffold-only controls at 2 and 8 weeks post implantation. Together, radiographs demonstrated accelerated healing of nonunion fractures in BMP2-expressing scaffolds. However, there was no significant difference observed between CG and CAG-MP. Notably, mineralization away from the cortex could be observed radiographically as early as week 6 (2-week after implantation) throughout the implanted scaffold surrounding the fracture site in cellular BMP2-LV-ASC scaffolds, indicating a potential risk of heterotopic ossification.

**Figure 4.**
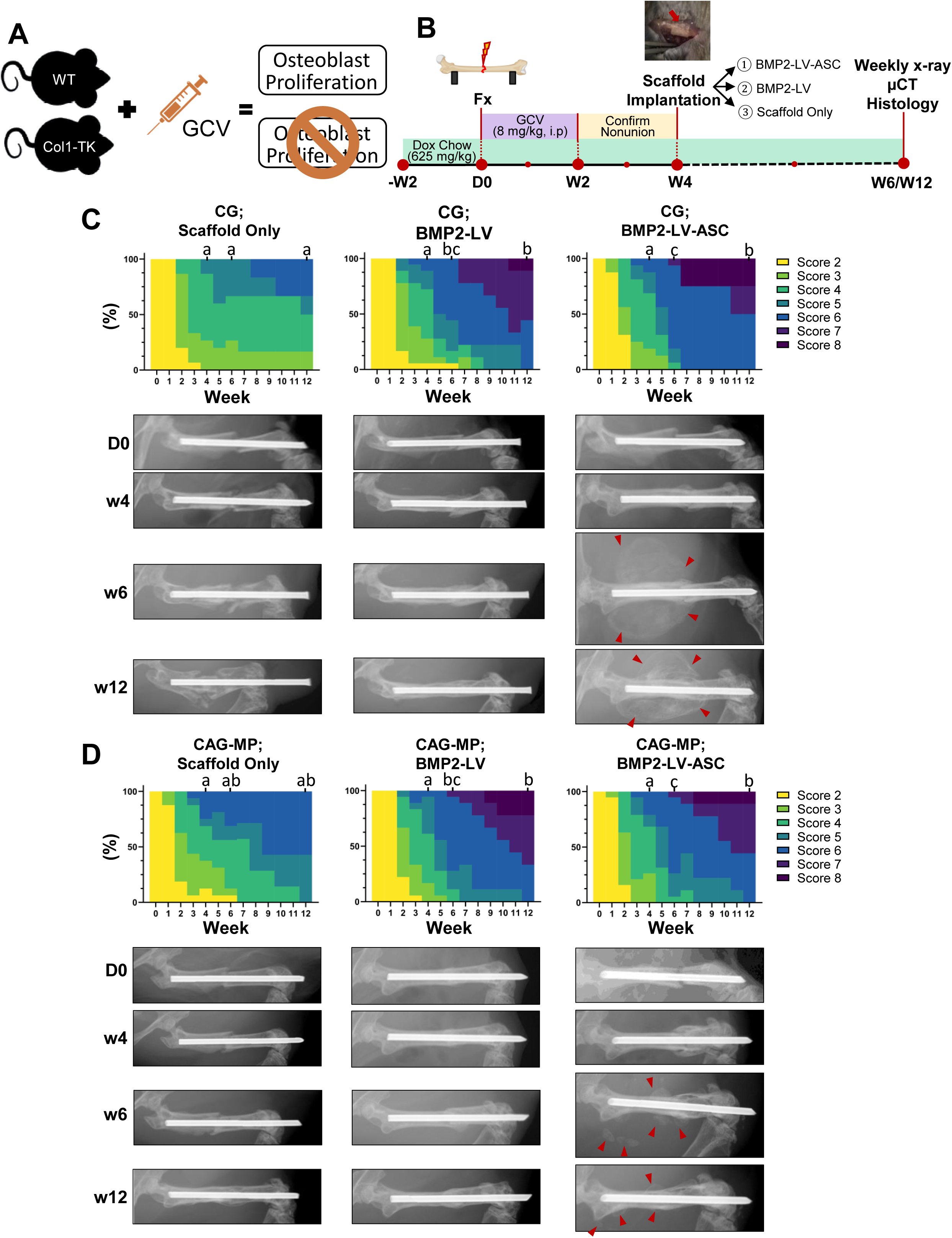
BMP2-expressing scaffolds promote radiographic healing in an atrophic nonunion fracture model. **(A)** Ganciclovir (GCV) administration to 3.6Col1a1-tk (Col-tk) mice ablates proliferating osteoblasts and producing atrophic nonunion; wild-type (WT) littermates retain normal osteoblast proliferation. **(B)** Experimental timeline. Femoral fracture was created at day 0, followed by GCV (8 mg/kg, i.p., once daily) for 2 weeks and doxycycline chow (625 mg/kg) until sacrifice. Nonunion was confirmed at week 4, at which point CG or CAG-MP scaffolds loaded with BMP2-LV-ASC, BMP2-LV, or left unmodified (scaffold only) were implanted at the fracture site. Radiographs were acquired weekly, with µCT and histology at week 6 or 12. **(C, D)** Weekly mRUST distributions and representative radiographs at day 0, week 4, 6, and 12 for **(C)** CG and **(D)** CAG-MP scaffolds. Stacked bars show the proportion of animals at each score; colors correspond to scores 2-8 as indicated. Red arrowheads indicate heterotopic mineralization (n =5-10/group). Groups sharing a letter are not significantly different (Kruskal-Wallis test); comparisons are made within each scaffold type across treatment groups at week 4, 6, and 12.

### BMP2-expressing Acellular Scaffolds (BMP2-LV) Induced Robust Callus Formation without Heterotopic Ossification

Healing was assessed using µCT at 2 and 8 weeks post-implantation. Callus volume was quantified by subtracting original cortical bone and marrow from total tissue volume, while heterotopic bone was defined as mineralized tissue not contiguous with the primary callus (Fig. 5, 6). At 2 weeks post-implantation, 3D reconstructions revealed persistent fracture lines in scaffold-only groups, whereas acellular BMP2-LV and cellular BMP2-LV-ASC groups displayed nearly complete callus bridging (Figs. 5A,B). Notably, heterotopic ossification was evident in cellular BMP2-LV-ASC groups. Quantitative analysis demonstrated that both BMP2-LV and BMP2-LV-ASC scaffolds had significantly increased callus bone volume (BV; Fig. 5C) and total volume (TV; Fig. 5D) compared with scaffold-only controls. Cellular BMP2-LV-ASC showed the highest mean BV and TV, but did not reach significance vs. acellular BMP2-LV. No significant differences were detected between the two scaffold types. Bone mineral density (BMD) was comparable among all groups (Fig. 5E), indicating that BMP2 primarily enhanced callus size rather than the density of bone within the callus. Notably, heterotopic bone formation (Fig. 5F) was pronounced in cellular BMP2-LV-ASC scaffolds but was absent (or negligible) in scaffold-only and acellular BMP2-LV groups. Notably, heterotopic bone formation was the only outcome that differed between scaffold types, being significantly greater in the CG group than in CAG-MP (p<0.0001).

**Figure 5.**
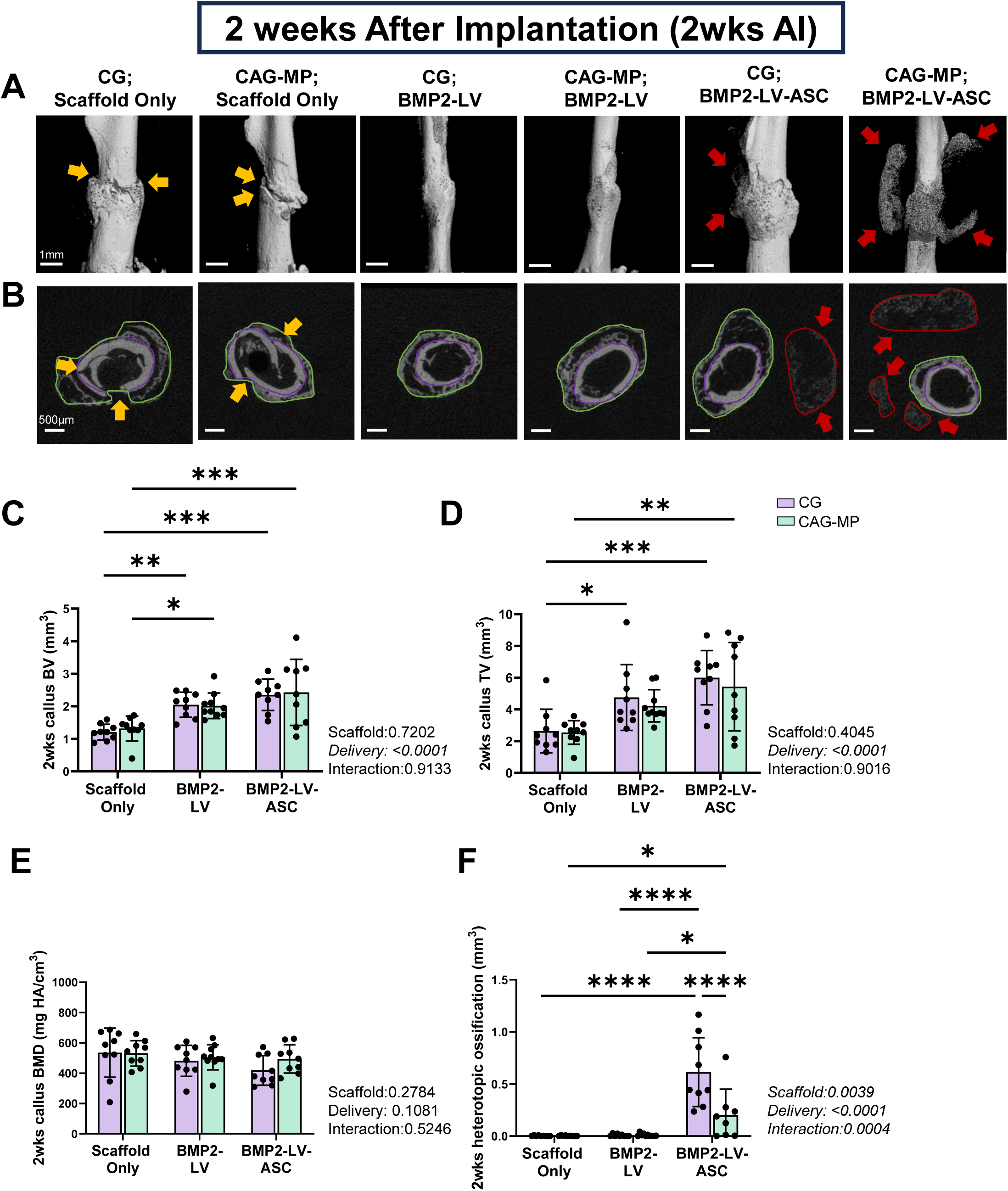
Micro-CT analysis of callus formation and heterotopic ossification at 2 weeks after implantation (2wks AI). **(A)** Representative µCT 3D reconstructions of fractured femurs 2 weeks after implantation of CG or CAG-MP scaffolds loaded with BMP2-LV-ASC, BMP2-LV, or left unmodified (scaffold only). Yellow arrowheads indicate unbridged fracture gaps; red arrowheads indicate heterotopic mineralization. Scale bar, 1mm. **(B)** Representative transverse µCT sections from the same specimens. Green outlines delineate total tissue volume, purple outlines the original cortical bone and marrow, and red outlines heterotopic mineralization, defined as mineralized tissue not contiguous with the primary callus. Scale bar, 500µm. **(C)** Callus bone volume (BV), **(D)** callus total tissue volume (TV), **(E)** callus bone mineral density (BMD), and **(F)** heterotopic mineralization. Callus volume was calculated by subtracting the original cortical bone and marrow from the total tissue volume. Data are mean ± SD, n=5-10/group. Two-way ANOVA with Tukey post-hoc comparisons; main effect and interaction p values are shown adjacent to each panel. *p < 0.05, **p < 0.01, ***p < 0.001, ****p < 0.0001.

At 8 weeks post-implantation, μCT 3D reconstructions demonstrated near-complete bone bridging in BMP2-treated groups compared to still visible fracture lines in the scaffold only groups (Fig. 6A). As determined by μCT analysis, both callus BV and TV were less at 8 weeks than at 2 weeks in all groups, reflecting ongoing bone remodeling and callus consolidation at the fracture site. Only CAG-MP BMP2-LV-ASC scaffolds maintained significantly higher callus BV than CAG-MP scaffold-only controls (1.38 ± 0.65mm³ vs. 0.72 ± 0.24mm³; p < 0.05; Fig. 6C). Similarly to 2 weeks, BMD remained comparable among all groups (Fig. 6E). However, the amount of heterotopic ossification at 8 weeks after implantation persisted in CAG-MP BMP2-LV-ASC group (mean = 0.27mm^3^; range = 0-0.96mm^3^, Fig. 6F) and was dramatically elevated in CG BMP2-LV-ASC group (mean = 5.73mm^3^; range = 0.02-16.47mm^3^; Fig. 6F), indicating exuberant osteogenesis in cellularized scaffolds. In summary, BMP2-expressing scaffolds promoted fracture healing at the fracture site, while acellular BMP2-LV scaffolds achieved substantial bone regeneration without heterotopic ossification.

**Figure 6.**
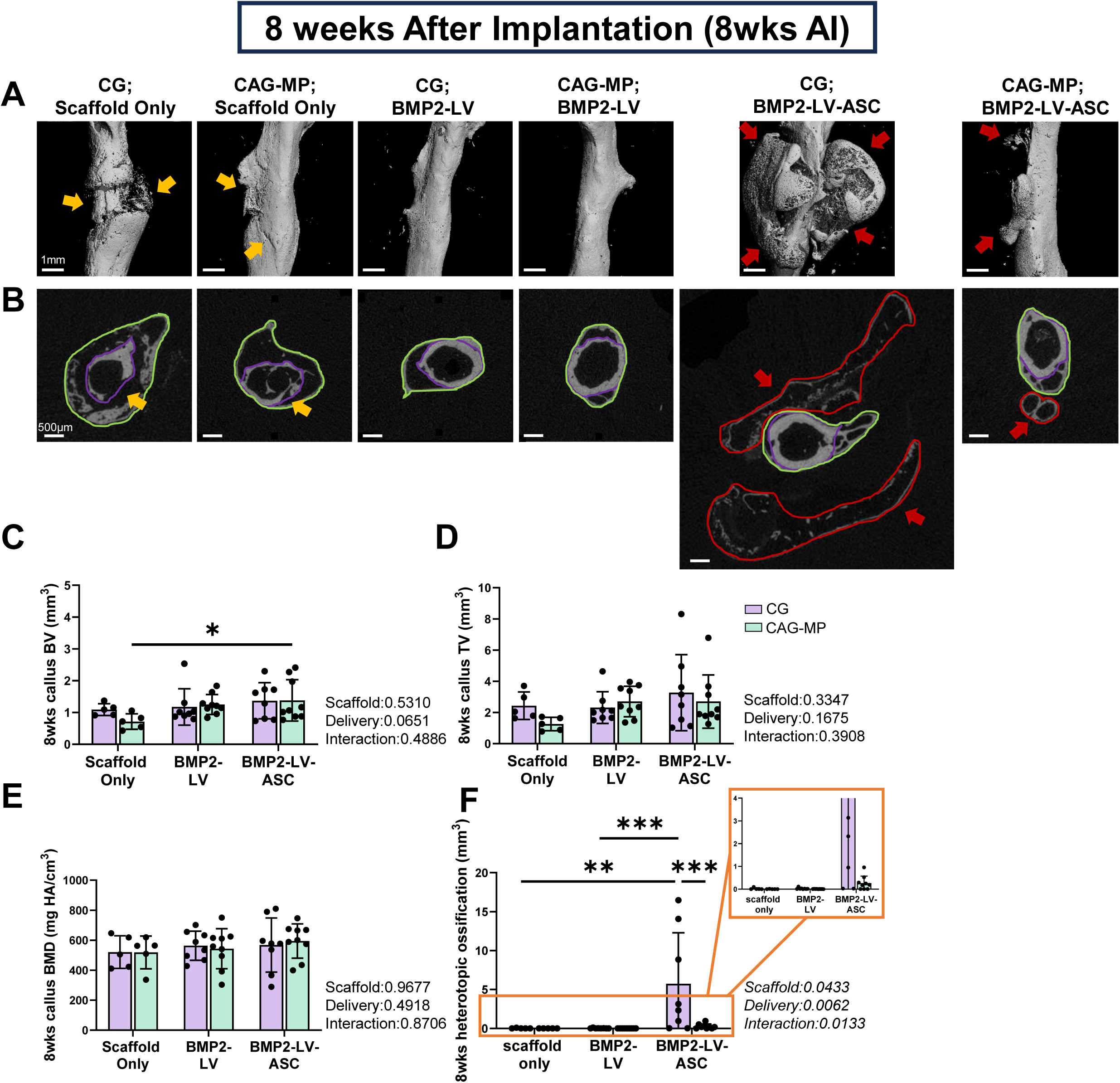
Micro-CT analysis of callus formation and heterotopic ossification at 8 weeks after implantation (8wks AI). **(A)** Representative µCT 3D reconstructions of fractured femurs 2 weeks after implantation of CG or CAG-MP scaffolds loaded with BMP2-LV-ASC, BMP2-LV, or left unmodified (scaffold only). Yellow arrowheads indicate unbridged fracture gaps; red arrowheads indicate heterotopic mineralization. Scale bar, 1mm. **(B)** Representative transverse µCT sections from the same specimens. Green outlines delineate total tissue volume, purple outlines the original cortical bone and marrow, and red outlines heterotopic mineralization, defined as not contiguous with the primary callus. Scale bar, 500µm. **(C)** Callus bone volume (BV), **(D)** callus total tissue volume (TV), **(E)** callus bone mineral density (BMD), and **(F)** heterotopic mineralization. Callus volume was calculated by subtracting the original cortical bone and marrow from the total tissue volume. Data are mean ± SD, n=5-10/group. Two-way ANOVA with Tukey post-hoc comparisons; main effect and interaction p values are shown adjacent to each panel. *p < 0.05, **p < 0.01, ***p < 0.001, ****p < 0.0001.

### BMP2-expressing Scaffolds Promoted Cortical Bridging and Accelerated Callus Maturation

Histological analysis of Picrosirius Red and Alcian Blue (PSRAB)-stained sections were performed at 2 and 8 weeks after implantation to further evaluate cortical bridging and assess callus tissue composition. At 2 weeks after implantation, fractures treated with scaffold only exhibited significantly smaller calluses compared with BMP2-expressing scaffolds (Fig. 7A, C), consistent with μCT findings. Complete cortical bridging was observed in 33% of scaffold-only samples, whereas 56% of cellular BMP2-LV-ASC scaffolds (both CG and CAG-MP) and 67% (CG) to 77.8% (CAG-MP) of acellular BMP2-LV scaffolds achieved full bridging (Fig. 7A). Notably, pronounced heterotopic ossification was observed in 89% CG BMP2-LV-ASC and 67% CAG-MP BMP2-LV-ASC samples. Quantification revealed that fractures treated with scaffold only contained significantly less woven bone and more fibrous tissue than those treated with BMP2-expressing scaffolds (Fig. 7A, Fig. 7B, D, E). Cartilage content was higher in the CAG-MP scaffold-only group than in the CAG-MP BMP2-LV group (Fig. 7A, B, F), while periosteal tissue content remained comparable among all groups (Fig. 7A, B, G). Interestingly, the only difference found between different scaffold types was in the cartilage content, where the CAG-MP; BMP2-LV-ASC group showed a trending higher cartilage content compared to the CG; BMP2-LV-ASC group. These findings indicated that BMP2-expressing scaffolds promoted callus mineralization at 2 weeks after implantation relative to scaffold-only controls.

**Figure 7.**
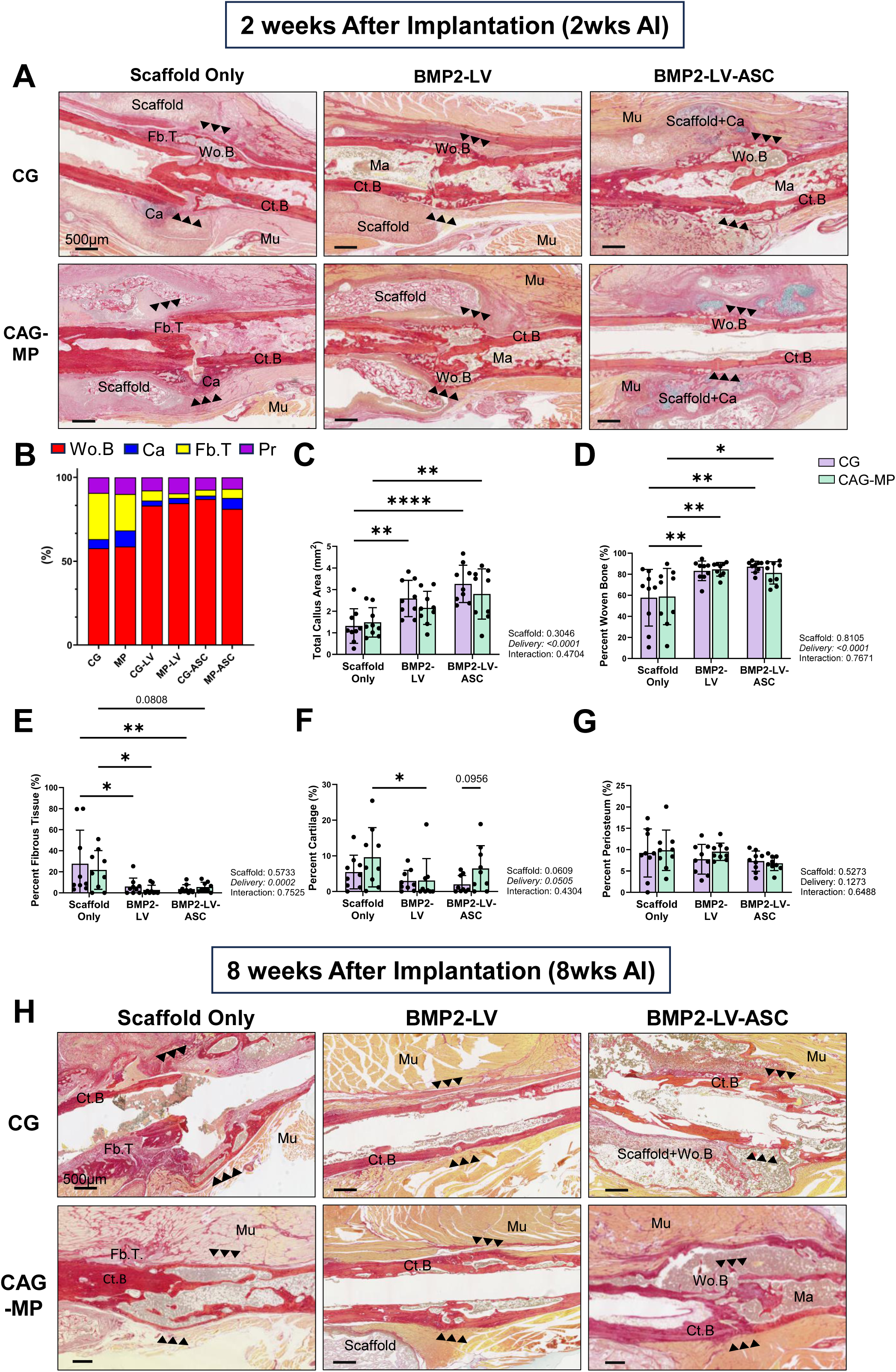
Histological analysis of callus composition at 2 (2wks AI) and 8 weeks after implantation (8wks AI). **(A)** Representative picrosirius red and alcian blue -stained (PSRAB) sections of fractured femurs 2 weeks after implantation (2wks AI) of CG or CAG-MP scaffolds loaded with BMP2-LV-ASC, BMP2-LV, or left unmodified (scaffold only). Ct.B, cortical bone; Wo.B, woven bone; Ca, cartilage; Fb.T, fibrous tissue; Ma, marrow; Mu, muscle; Ps, periosteum. Arrowheads indicate the fracture site. Scale bars, 500µm. **(B)** Relative tissue composition of the callus at two weeks, expressed as percentage of total callus area. **(C)** Total callus area, **(D)** percent woven bone, **(E)** percent fibrous tissue, **(F)** percent cartilage, and **(G)** percent periosteum at two weeks after implantation. **(H)** Representative sections at 8 weeks post-implantation, labeled as in (A). Scale bars, 500µm. Data are mean ± SD, n=5-10/group. Two-way ANOVA with Tukey post-hoc comparisons; main effect and interaction p values are shown adjacent to each panel. *p < 0.05, **p < 0.01, ****p < 0.0001;

At 8 weeks after implantation, histology demonstrated complete cortical bridging in all BMP2-LV (both CG and CAG-MP) and CAG-MP; BMP2-LV-ASC samples (Fig. 7H). In contrast, only 50% of CG and 57% of CAG-MP scaffold-only samples achieved full bridging. Two-thirds (67%) of CAG-MP BMP2-LV-ASC samples exhibited heterotopic bone formation, while 75% of CG BMP2-LV-ASC samples had heterotopic bone formation, and only 50% showed full bridging. Furthermore, fully bridged cortices without a visible fracture line were seen in 33% of CG and 30% of CAG-MP; BMP2-LV samples, as well as 12.5% of CG and 33% CAG-MP BMP2-LV-ASC (Fig. 7H), indicating full healing with remodeling of the fracture site. In addition, the callus as well as the heterotopic tissue were mostly mineralized regardless of callus size or bridging condition among all groups (data not shown), indicating that the completion of the remodeling phase at this time point. In summary, acellular and cellular BMP2-expressing scaffolds accelerated fracture healing and promoted cortical bridging in atrophic nonunion fractures. However, acellular scaffolds (BMP2-LV) did not stimulate heterotopic mineralization in contrast to BMP2-LV-ASC scaffolds.

### 3-day Doxycycline Induction Preserves Callus Formation while Reducing Heterotopic Ossification

To possibly mitigate the severe heterotopic ossification observed in BMP2-LV-ASC groups, we withdrew doxycycline 3 days after scaffold implantation, limiting transgene expression to a three-day window (Fig. 8A). Because heterotopic ossification was most pronounced in CG BMP2-LV-ASC constructs (Fig. 5F, 6F), the shortened induction was evaluated only in this group. Compared to CG BMP2-LV-ASC treated with continuous doxycycline (ASC_cont), 3-day induction (ASC_3d) markedly reduced heterotopic ossification while preserving callus formation (Fig. 8B, C). µCT revealed comparable BV (Fig. 8D,H), TV (Fig. 8E, I), and BV/TV (Fig. 8F, J) between ASC_cont and ASC_3d groups at 2 weeks after implantation. At 8 weeks, BV was significantly higher in the ASC_3d group, while TV and BV/TV did not differ between groups. We also noted that BV/TV was higher in CG scaffold only group than ASC_3d at 2 weeks after implantation, although this is likely reflecting the small callus volume (TV) in the CG scaffold only group. At 2 weeks after implantation, the heterotopic ossification in ASC_3d remained greater than in CG scaffold only and CG BMP2-LV (LV_cont) groups, but was substantially lower than in ASC_cont (Fig. 8G). By 8 weeks after implantation, heterotopic ossification in ASC_3d group was negligible and no longer differed from either scaffold only or LV_cont (Fig. 8K). Similarly, histological analysis exhibited comparable callus composition between ASC_3d and the ASC_cont and LV_cont groups at 2 weeks post implantation (Fig. 9A, B), including total callus area (Fig. 9C) and woven bone composition (Fig. 9D), both of which exceeded scaffold-only controls. Together, these results indicate that limiting doxycycline induction to three days constrains BMP2 expression sufficiently to prevent heterotopic ossification while preserving the osteogenic response.

**Figure 8.**
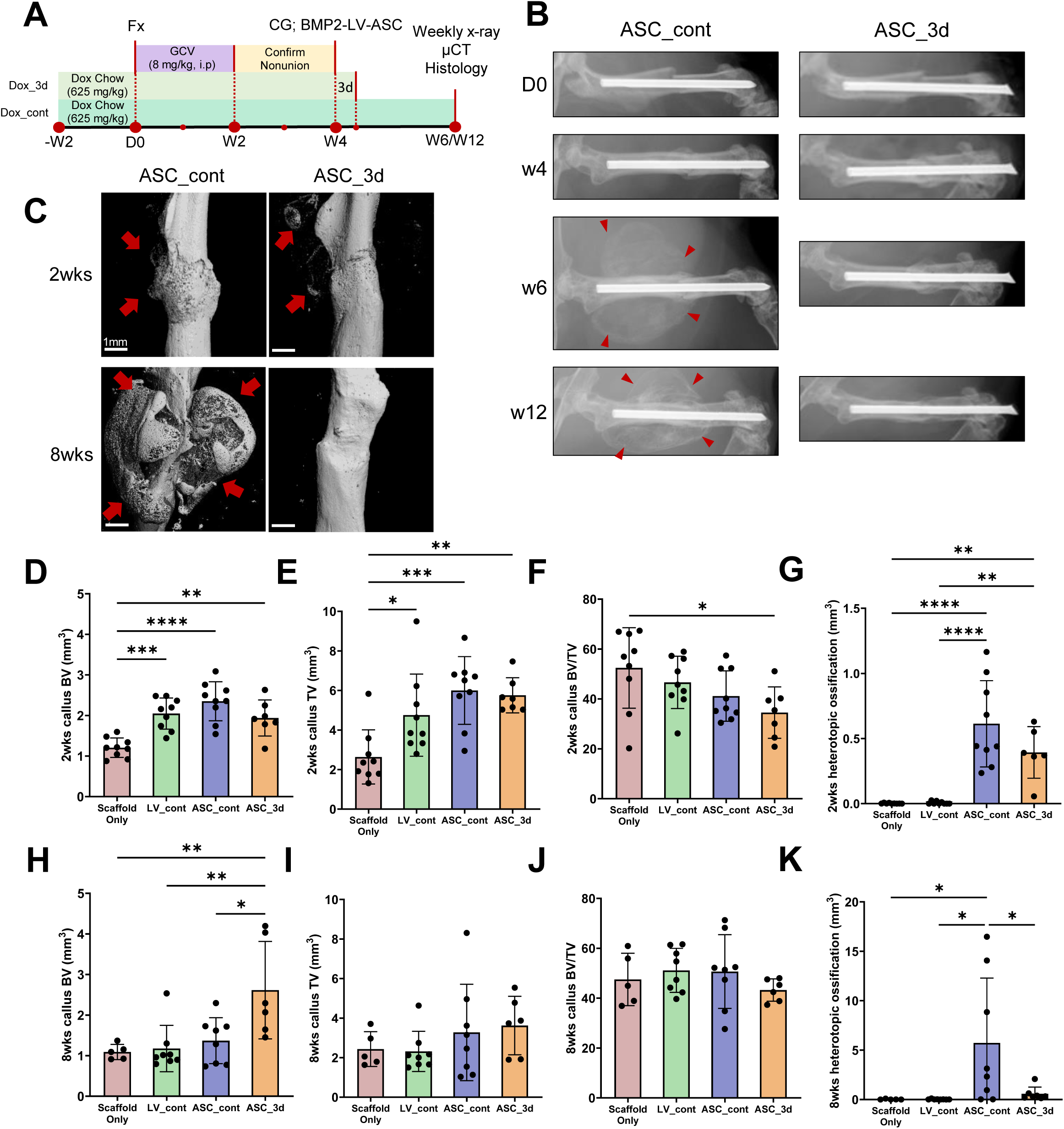
Three-day induction of BMP2 preserves callus formation while reducing heterotopic ossification. **(A)** Experimental timeline. Femoral fracture was created at day 0, followed by ganciclovir (8 mg/kg, i.p., once daily) for 2 weeks. Nonunion was confirmed at week 4, at which point CG scaffolds loaded with BMP2-LV-ASC were implanted. Mice received doxycycline chow (625mg/kg) either continuously from day 0 until sacrifice (Dox_cont) or for 3 days following implantation (Dox_3d). Radiographs were acquired weekly, with µCT and histology at week 6 or 12. **(B)** Representative radiographs at days 0, 4, 6, and 12 weeks. Red arrowheads indicate heterotopic mineralization. **(C)** Representative µCT 3D reconstructions at 2 and 8 weeks after implantation. Red arrowheads indicate ectopic mineralization. Scale bars, 1mm. **(D-G)** Callus bone volume (BV), callus total tissue volume (TV), callus BV/TV, and heterotopic ossification volume at 2 weeks. **(H-K)** The corresponding measurements at 8 weeks. Data are mean ± SD, n =5-10/group. One-way ANOVA with Tukey’s post-hoc test. *p < 0.05, **p < 0.01, ***p < 0.001, ****p < 0.0001. LV_cont: BMP2-LV with continuous doxycycline; ASC_cont, BMP2-LV-ASC with continuous doxycycline; ASC_3d, BMP2-LV-ASC with three-day doxycycline. Values for the scaffold-only, LV_cont, and ASC_cont groups are the CG data reported in Fig. 5&6, included here as reference comparators.

**Figure 9.**
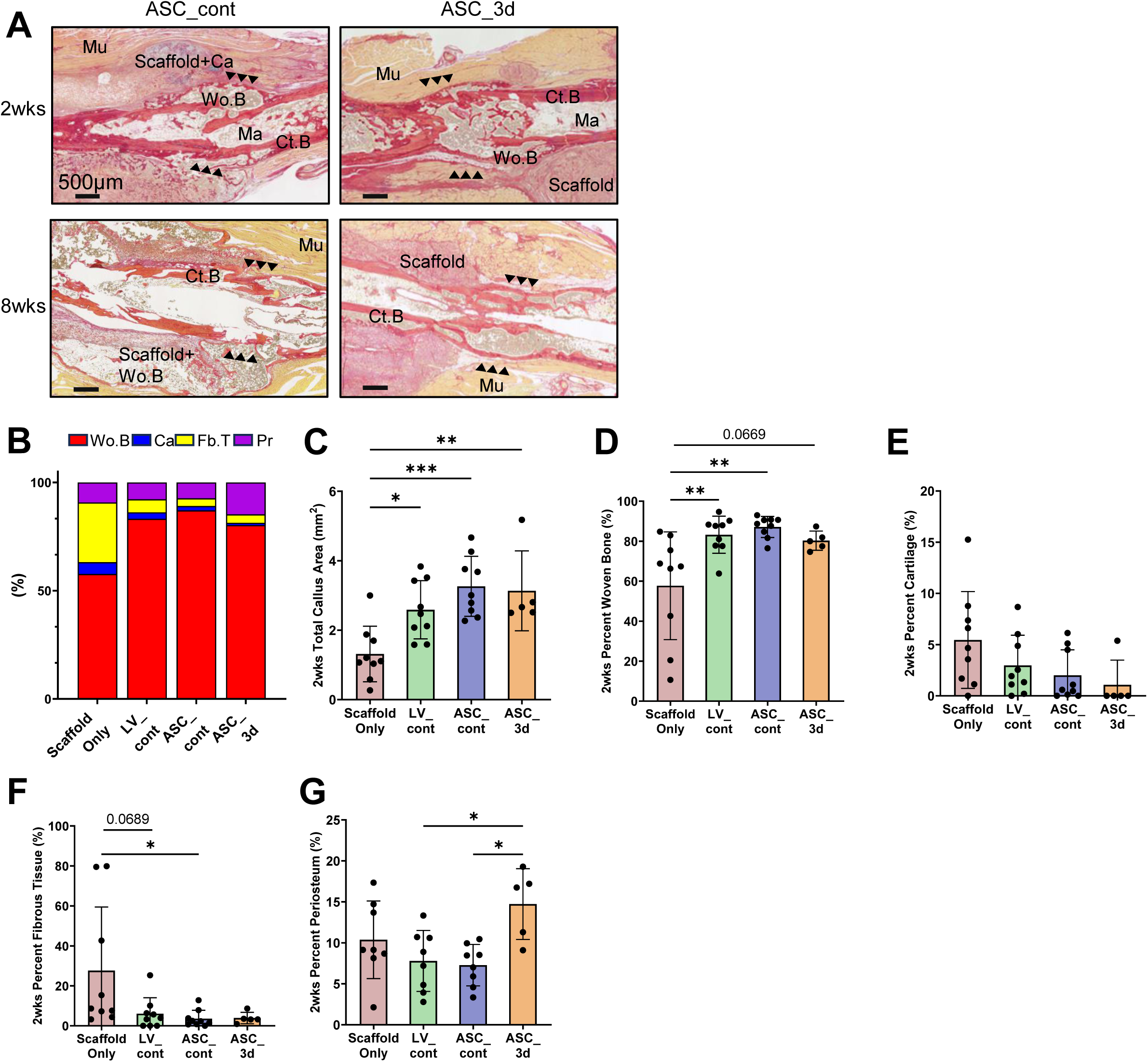
Histological analysis of callus composition following continuous or 3-day BMP2 induction. **(A)** Representative picrosirius red and alcian blue-stained (PSRAB) sections of fractured femurs at 2 and 8 weeks after implantation of CG BMP2-LV-ASC scaffolds, with doxycycline administered continuously (ASC_cont) or from day 0 to 3 days following implantation (ASC_3d). Ct.B, cortical bone; Wo.B, woven bone; Ca, cartilage; Ma, marrow; Mu, muscle. Arrowheads indicate the fracture site. Scale bars, 500µm. **(B)** Relative tissue composition of the callus at 2 weeks after implantation, expressed as percentage of total callus area. Wo.B, woven bone; Ca, cartilage; Fb.T, fibrous tissue; Pr, periosteum. **(C)** Total callus area, **(D)** percent woven bone, **(E)** percent cartilage, **(F)** percent fibrous tissue, and **(G)** percent periosteum at 2 weeks. LV_cont: BMP2-LV with continuous doxycycline; ASC_cont, BMP2-LV-ASC with continuous doxycycline; ASC_3d, BMP2-LV-ASC with 3-day doxycycline. Data are mean ± SD, n=5-10/group. One-way ANOVA with Tukey’s post-hoc test. *p < 0.05, **p < 0.01, ***p < 0.001. Values for the scaffold-only, LV_cont, and ASC_cont groups are the CG data reported in Fig. 7, included here as reference comparators.

## Discussion

We hypothesized that by incorporating agarose in chitosan-gelatin cryogel scaffolds, the pore size and porosity could be increased, thereby improving cell infiltration and mechanical performance. Compared to CG scaffolds, CAG scaffolds exhibited larger pore size, higher porosity and greater cell infiltration *in vivo*. Despite having a lower modulus, CAG scaffolds showed a shift towards more ductile failure with increased post-yield toughness. We further hypothesized that localized delivery of doxycycline-inducible BMP2-LV via these scaffolds would rescue atrophic nonunion while allowing temporal control of transgene expression. *In vivo*, both cellular (BMP2-LV-ASC) and acellular (BMP2-LV) scaffolds receiving continuous doxycycline induction (Dox_cont) rescued atrophic nonunion, indicated by increased cortical bridging and accelerated callus maturation. However, cellular delivery produced substantial heterotopic ossification adjacent to the fracture site, which was especially severe in CG scaffolds. By comparison, acellular scaffolds achieved comparable fracture healing while inducing minimal heterotopic ossification. When the administration of doxycycline was restricted to only 3 days after scaffold implantation (Dox_3d), heterotopic ossification was largely eliminated in cellular CG scaffolds while callus formation was maintained. This result is consistent with the doxycycline-dependent BMP2 secretion we observed *in vitro*, although we note that for *in vitro* BMP2 production, ST2 cells were used, instead of ASCs.

The murine model that we used in this study is driven by the ganciclovir-mediated ablation of proliferating osteoblasts, which prevents callus formation and results in a clinically relevant atrophic nonunion[11]. By delivering BMP2 using cryogel scaffolds, both acellular (BMP2-LV) and cellular (BMP2-LV-ASC) constructs promoted cortical bridging and increased callus bone volume relative to scaffold-only controls. Callus bone mineral density did not differ between groups, indicating that BMP2 expression increased the quantity of bone formed rather than the density of the callus tissue. Histological analysis showed similar results, where BMP2-expressing scaffolds resulted in calluses with significantly more woven bone and less fibrous tissue than controls, indicating progression beyond the fibrous, atrophic state that characterizes this model if left untreated. By 8 weeks, the callus in scaffold-treated groups was completely bony and callus bone volume was similar to control groups, which is consistent with remodeling of the treated callus[36,37]. Together, these findings extend previous work showing that scaffold-mediated gene delivery supports sustained transgene expression and protein synthesis by MSCs[15,25,38], and are consistent with the sustained BMP2 secretion observed *in vitro* through day 10. Importantly, acellular and cellular deliveries produced comparable healing, indicating that transduction of host cells at the defect site was sufficient to drive repair. To our knowledge, cellularized and acellular delivery of lentivirus have not previously been compared in a single study. Our results show that *in vivo* transduction is sufficient to rescue nonunion healing with efficacy comparable to cellular delivery.

One important difference between the two delivery strategies is that, under sustained doxycycline induction, heterotopic ossification was prevalent with cellular but not acellular scaffolds. This was observed adjacent to the fracture site, where heterotopic bone formed in the surrounding soft tissue area, separate from the callus bone. We previously reported similar heterotopic ossification in cellularized CG scaffolds delivering BMP2-LV[15]. This phenomenon recapitulates a known liability of BMP2-based therapeutics. Despite its effectiveness in promoting spinal fusion clinically, the supraphysiologic doses required for efficacy with rhBMP-2 have been associated with ectopic bone formation and other dose-related complications[19,20]. Development of novel delivery strategies that constrain both the magnitude and the location of BMP2 activity are therefore warranted. We attribute the formation of heterotopic ossification to the quantity and spatial distribution of BMP2-producing cells rather than to the vector itself. In cellular delivery, ASCs were transduced *ex vivo* as a large population and ready to express transgene upon implantation. It is possible that these transduced ASCs migrate from the scaffold to the surrounding soft tissues, resulting in BMP2 expression beyond the intended site. By contrast, in acellular delivery, BMP2 production requires transduction of host cells that migrate into the scaffolds, and therefore transgene expression was likely more gradual and localized to the fracture site. However, we note that expression of *Bmp2* in the subcutaneously implanted scaffolds was similar between acellular BMP2-LV and cellularized BMP2-LV-ASC constructs. These measurements were restricted to the scaffolds and therefore would not capture expression by cells that might have migrated into the surrounding muscle. Heterotopic and orthotopic sites are substantially different regarding progenitor populations, cytokine milieu, and vascular supply, and identical constructs have been shown to produce different amounts of bone at the two locations in mice despite equivalent cell survival[39,40]. Thus, expression within the scaffold alone provides an incomplete account of the heterotopic bone response.

To mitigate the heterotopic ossification caused continued induction of BMP2 production by BMP2-LV-ASC constructs, we shortened doxycycline administration to three days following scaffold implantation to limit BMP2 secretion. µCT and histological analysis showed that CG BMP2-LV-ASC animals receiving three-day induction (ASC_3d) developed significantly less heterotopic ossification than those maintained on continuous doxycycline (ASC_cont), while callus bone volume did not differ between groups at 2 weeks and was significantly greater in ASC_3d at 8 weeks. This degree of control is not achievable with rhBMP2 protein delivery. With a half-life of ∼10 minutes[41], rhBMP2 must be delivered at supraphysiologic doses from carriers whose release profile is fixed at implantation[42]. By contrast, transduced cells produce BMP2 continuously. *In vitro* ELISA confirmed that BMP2 secretion continuously increases to day 6 and maintains through day 10 with doxycycline induction. Since the secretion required both lentivirus and doxycycline, the duration of production could be altered after implantation simply by withdrawing doxycycline from the diet. Endogenous BMP2 expression during fracture healing is itself not constant, rising shortly after injury, falling transiently, and increasing again during mineralization of the cartilaginous callus before returning to baseline[42–44]. A regulated vector therefore offers the possibility of approximating this temporal profile rather than imposing continuous expression, which may serve as a more effective strategy for avoiding the adverse effects of supraphysiologic rhBMP2 protein delivery.

The incorporation of agarose changed the pore architecture and mechanical behavior of the cryogel scaffolds, but these differences did not translate into differences in fracture healing. Moreover, we were able to increase pore size in the CAG series by slowing the freezing rate[45]. Overall, the pore size and porosity of CG and CAG scaffolds were comparable to previously reported values for similar formulations[29,30]. Mechanically, CAG-SP exhibited a lower modulus but greater ultimate strain and post-yield toughness than CG, shifting failure behavior from brittle to ductile which made CAG scaffolds more robust for surgical manipulation. Cell infiltration increased nonlinearly with pore size; although CAG-LP had the largest pores, infiltration was greatest in CAG-MP, suggesting that intermediate pore sizes balance accessibility against the surface area available for cell attachment [46]. Despite these structural and mechanical differences, callus formation and radiographic healing were comparable between CG and CAG-MP. Scaffold composition did influence the surrounding tissue response to BMP2 transgene, as CG BMP2-LV-ASC constructs produced significantly more heterotopic ossification than CAG-MP. This might be due to differences in crosslink chemistry. CG scaffolds were crosslinked with 1% glutaraldehyde compared with 0.2% for CAG, and glutaraldehyde-fixed biomaterials are well documented to undergo dystrophic calcification *in vivo*, particularly in subcutaneous implantation models, an effect attributed largely to residual free aldehyde groups rather than to crosslink density itself[47,48]. Regional mineral deposition will elevate the extracellular concentration of calcium ions at the implantation site. In the absence of vector, this can induce macrophages and mesenchymal cells to express and secrete BMP2[47,48] and raise baseline *Bmp2* modestly while amplifying the osteogenic response once BMP2 is supplied[49]. However, the relationship between glutaraldehyde concentration and calcification is not straightforward; some studies reported that with higher fixative concentrations, calcification was reduced in some ranges[50]. Differences in vector retention and release kinetics[51], as well as substrate stiffness[52] effects on osteogenic differentiation may also contribute to differences between scaffold groups. Because CG and CAG differ in both crosslinker concentration and agarose content, and therefore in porosity, pore size, stiffness, and degradation behavior, we cannot identify a single cause of the scaffold effect on heterotopic ossification.

Atrophic nonunion remains among the most difficult problems in fracture care. Despite the prevalence of nonunions, even in the absence of bone loss, most pre-clinical studies of bone repair focus on segmental, critical-sized defects. Clinically, these are two different scenarios and may require different treatment approaches. The current standard for nonunion is surgical debridement and hardware revision, with outcomes unpredictable[5,6]. The acellular lentiviral approach we evaluated transduces host cells *in vivo*, such that no autologous bone graft or cell harvest, *ex vivo* expansion, or GMP cell manufacturing is required. Both the scaffold and the vector can be manufactured and stored in advance and combined shortly before surgery, avoiding the weeks of patient-specific cell culture required for autologous cell therapy; the construct could be implanted during a revision procedure once nonunion is confirmed. rhBMP-2 protein is potently osteoinductive, but once a carrier is implanted, neither the dose nor the duration of release can be changed[19,20]. By comparison, a doxycycline-dependent lentiviral vector allows expression to be controlled in duration, permitting treatment to be stopped before heterotopic bone or other adverse responses arise. Moreover, doxycycline is itself a widely used approved agent with well-characterized pharmacokinetics[53]. Nonetheless, substantial barriers to translation still need to be considered, as lentiviral vectors integrate permanently into the host genome, carrying a theoretical risk of insertional mutagenesis[21].

Several limitations should be noted. The 3-day doxycycline regimen was tested only in CG BMP2-LV-ASC constructs, so it remains unknown whether shortened induction would produce comparable effects in CAG scaffolds or in acellular constructs. Doxycycline has been reported to suppress osteoclast activity and increase bone formation in rodents[54,55]. This effect would be expected to slow remodeling in the continuously treated animals leading to a larger callus, yet callus bone volume at 8 weeks was greater with 3-day induction, making it unlikely that there was a direct pharmacological effect of doxycycline. A vector-free doxycycline-duration control group would nonetheless be required to confirm this. In this study, we only tested a single shortened induction window. Future study will be required to define the relationship between induction timing, duration, callus healing and heterotopic bone formation, including induction aligned to specific stages of healing. Several measurements were also limited in scope. *Bmp2* expression was quantified only in scaffold-associated tissue in the subcutaneous model. Therefore, the expression by cells that had migrated into surrounding soft tissue, and expression at the fracture site itself, were not captured. Lentiviral transduction *in vivo* was characterized qualitatively by confocal microscopy in the subcutaneous model rather than quantified at the fracture site. CAG pore-size variants were pooled for qPCR and subcutaneous mineralization analyses because differences among them were negligible, which limits the resolution of these comparisons with respect to pore architecture. In addition, tensile testing was only performed with CAG-SP scaffolds, instead of CAG-MP scaffolds. With higher porosity, CAG-MP should exhibit higher post yield toughness and ultimate strain than CAG-SP scaffolds, but future study will still be required to evaluate the mechanical properties of CAG-MP directly. Finally, functional outcomes including animal behavior and biomechanical properties of the healed femur were not evaluated; both are important variables to assess in future studies.

In summary, delivery of doxycycline inducible BMP2-lentivirus via chitosan-based cryogel scaffolds effectively rescued fracture healing in a clinically relevant, murine nonunion model. Acellular delivery produced comparable healing compared to BMP2-transduced ASCs with substantially less heterotopic ossification, which indicates that transduction of the host cells at the fracture site is sufficient to heal atrophic nonunion. When cells were used, restricting doxycycline induction of BMP2 to a short period after implantation significantly reduced heterotopic ossification. This latter finding indicates that the timing of BMP2 production, rather than the total dosage, is an important variable in promoting healing while reducing adverse effects. With an inducible vector, the local protein expression can be adjusted non-invasively, offering a means to match BMP2 delivery to the temporal profile of endogenous expression that occurs during normal healing.

## Data availability

Data will be made available on request.

## Funding Acknowledgement

DOD PW81XWH-21-1-0386 (to MJS); NIH P30 AR074992; NIH S10 OD028573; NIH AR080902 (to FG); NIH NCRR 1S10RR027552; The views expressed are those of the authors and do not necessarily represent the official views of the National Institutes of Health.

## References

[1] B. Wildemann, A. Ignatius, F. Leung, L.A. Taitsman, R.M. Smith, R. Pesántez, M.J. Stoddart, R.G. Richards, J.B. Jupiter, Non-union bone fractures, Nat Rev Dis Primers 7 (2021) 57. 10.1038/s41572-021-00289-8.

[2] M.F. Vanderkarr, J.W. Ruppenkamp, M. Vanderkarr, C.E. Holy, M. Blauth, Risk factors and healthcare costs associated with long bone fracture non-union: a retrospective US claims database analysis, J Orthop Surg Res 18 (2023) 745. 10.1186/s13018-023-04232-3.

[3] J. Nicholson, N. Makaram, A. Simpson, J. Keating, Fracture nonunion in long bones: A literature review of risk factors and surgical management, Injury 52 (2021) S3–S11. 10.1016/j.injury.2020.11.029.

[4] R. Dimitriou, E. Jones, D. McGonagle, P.V. Giannoudis, Bone regeneration: current concepts and future directions, BMC Med 9 (2011) 66. 10.1186/1741-7015-9-66.

[5] T.T. Roberts, A.J. Rosenbaum, Bone grafts, bone substitutes and orthobiologics: The bridge between basic science and clinical advancements in fracture healing, Organogenesis 8 (2012) 114–124. 10.4161/org.23306.

[6] H.-S. Sohn, J.-K. Oh, Review of bone graft and bone substitutes with an emphasis on fracture surgeries, Biomater Res 23 (2019) 9. 10.1186/s40824-019-0157-y.

[7] T. Tawonsawatruk, M. Kelly, H. Simpson, Evaluation of Native Mesenchymal Stem Cells from Bone Marrow and Local Tissue in an Atrophic Nonunion Model, Tissue Engineering Part C: Methods 20 (2014) 524–532. 10.1089/ten.tec.2013.0465.

[8] M.I. Pérez Núñez, D. Ferreño Blanco, A. Alfonso Fernández, J.A. Casado De Prado, M. Sánchez Crespo, M. De La Red Gallego, A. Pascual Carra, T. Rodriguez López, S. Diego Cavia, C. Garcés Zarzalejo, M. Mayorga Fernández, E. Ruiz Martínez, I. Carrascal Vaquero, J.A. Riancho Moral, Comparative study of the effect of PTH (1–84) and strontium ranelate in an experimental model of atrophic nonunion, Injury 46 (2015) 2359–2367. 10.1016/j.injury.2015.10.007.

[9] R.R. Kelly, M.A. McCrackin, D.L. Russell, L.R. Leddy, J.J. Cray, A.C. LaRue, Murine Aseptic Surgical Model of Femoral Atrophic Nonunion, MethodsX 7 (2020) 100898. 10.1016/j.mex.2020.100898.

[10] E.A. Lin, C.-J. Liu, A. Monroy, S. Khurana, K.A. Egol, Prevention of Atrophic Nonunion by the Systemic Administration of Parathyroid Hormone (PTH 1–34) in an Experimental Animal Model, Journal of Orthopaedic Trauma 26 (2012) 719–723. 10.1097/BOT.0b013e31826f5b9e.

[11] K.R. Hixon, J.A. McKenzie, D.A.W. Sykes, S. Yoneda, A. Hensley, E.G. Buettmann, H. Zheng, D. Skouteris, A. McAlinden, A.N. Miller, M.J. Silva, Ablation of Proliferating Osteoblast Lineage Cells After Fracture Leads to Atrophic Nonunion in a Mouse Model, Journal of Bone and Mineral Research 36 (2021) 2243–2257. 10.1002/jbmr.4424.

[12] R.L. Jilka, C.A. O’Brien, A.A. Ali, P.K. Roberson, R.S. Weinstein, S.C. Manolagas, Intermittent PTH stimulates periosteal bone formation by actions on post-mitotic preosteoblasts, Bone 44 (2009) 275–286. 10.1016/j.bone.2008.10.037.

[13] A. Kavitha Sri, C. Arthi, N.R. Neya, G.S. Hikku, Nano-hydroxyapatite/collagen composite as scaffold material for bone regeneration, Biomed. Mater. 18 (2023) 032002. 10.1088/1748-605X/acc99e.

[14] K.R. Hixon, T. Lu, S.A. Sell, A comprehensive review of cryogels and their roles in tissue engineering applications, Acta Biomaterialia 62 (2017) 29–41. 10.1016/j.actbio.2017.08.033.

[15] K.R. Hixon, D.B. Katz, J.A. McKenzie, A.N. Miller, F. Guilak, M.J. Silva, Cryogel Scaffold-Mediated Delivery of Adipose-Derived Stem Cells Promotes Healing in Murine Model of Atrophic Non-Union, Front. Bioeng. Biotechnol. 10 (2022) 851904. 10.3389/fbioe.2022.851904.

[16] X. Wang, Z. Yuan, G. Cai, Y. Lu, S. Zhou, P. Shang, C. Li, Z. Wang, Z. Liu, M. Shafiq, M. El-Newehy, M.M. Abdulhameed, Xiao Lu, Y. Xu, X. Mo, Flexible Short Silica Fibers and Tricalcium Phosphate Synergistically Promote Bone Fracture Healing in Composite Cryogel Scaffolds, Advanced Healthcare Materials 14 (2025). 10.1002/adhm.202404329.

[17] B.Y. Ozturk, I. Inci, S. Egri, A.M. Ozturk, H. Yetkin, G. Goktas, C. Elmas, E. Piskin, D. Erdogan, The treatment of segmental bone defects in rabbit tibiae with vascular endothelial growth factor (VEGF)-loaded gelatin/hydroxyapatite “cryogel” scaffold, Eur J Orthop Surg Traumatol 23 (2013) 767–774. 10.1007/s00590-012-1070-4.

[18] C.E. Gillman, A.C. Jayasuriya, FDA-approved bone grafts and bone graft substitute devices in bone regeneration, Materials Science and Engineering: C 130 (2021) 112466. 10.1016/j.msec.2021.112466.

[19] S.-Y. Park, K.-H. Kim, S. Kim, Y.-M. Lee, Y.-J. Seol, BMP-2 Gene Delivery-Based Bone Regeneration in Dentistry, Pharmaceutics 11 (2019) 393. 10.3390/pharmaceutics11080393.

[20] E.J. Carragee, E.L. Hurwitz, B.K. Weiner, A critical review of recombinant human bone morphogenetic protein-2 trials in spinal surgery: emerging safety concerns and lessons learned, The Spine Journal 11 (2011) 471–491. 10.1016/j.spinee.2011.04.023.

[21] X. Li, Y. Le, Z. Zhang, X. Nian, B. Liu, X. Yang, Viral Vector-Based Gene Therapy, International Journal of Molecular Sciences 24 (2023). 10.3390/ijms24097736.

[22] E. Abordo-Adesida, A. Follenzi, C. Barcia, S. Sciascia, M.G. Castro, L. Naldini, P.R. Lowenstein, Stability of Lentiviral Vector-Mediated Transgene Expression in the Brain in the Presence of Systemic Antivector Immune Responses, Human Gene Therapy 16 (2005) 741–751. 10.1089/hum.2005.16.741.

[23] J.M. Brunger, N.P.T. Huynh, C.M. Guenther, P. Perez-Pinera, F.T. Moutos, J. Sanchez-Adams, C.A. Gersbach, F. Guilak, Scaffold-mediated lentiviral transduction for functional tissue engineering of cartilage, Proc. Natl. Acad. Sci. U.S.A. 111 (2014). 10.1073/pnas.1321744111.

[24] K.A. Glass, J.M. Link, J.M. Brunger, F.T. Moutos, C.A. Gersbach, F. Guilak, Tissue-engineered cartilage with inducible and tunable immunomodulatory properties, Biomaterials 35 (2014) 5921–5931. 10.1016/j.biomaterials.2014.03.073.

[25] C.R. Rowland, K.A. Glass, A.R. Ettyreddy, C.C. Gloss, J.R.L. Matthews, N.P.T. Huynh, F. Guilak, Regulation of decellularized tissue remodeling via scaffold-mediated lentiviral delivery in anatomically-shaped osteochondral constructs, Biomaterials 177 (2018) 161–175. 10.1016/j.biomaterials.2018.04.049.

[26] B. Wang, Y. Guo, X. Chen, C. Zeng, Q. Hu, W. Yin, W. Li, H. Xie, B. Zhang, X. Huang, F. Yu, Nanoparticle-modified chitosan-agarose-gelatin scaffold for sustained release of SDF-1 and BMP-2, IJN Volume 13 (2018) 7395–7408. 10.2147/IJN.S180859.

[27] E. Sartip, T. Behzad, M. Kharaziha, Fabrication and characterization of 3D printed agarose/poly(ethylene glycol) diacrylate/hydroxyapatite nanocomposite hydrogel for cartilage tissue engineering, International Journal of Biological Macromolecules 320 (2025) 145573. 10.1016/j.ijbiomac.2025.145573.

[28] F.T. Moutos, K.A. Glass, S.A. Compton, A.K. Ross, C.A. Gersbach, F. Guilak, B.T. Estes, Anatomically shaped tissue-engineered cartilage with tunable and inducible anticytokine delivery for biological joint resurfacing, Proc Natl Acad Sci U S A 113 (2016) E4513–4522. 10.1073/pnas.1601639113.

[29] S. Bhat, A. Tripathi, A. Kumar, Supermacroprous chitosan–agarose–gelatin cryogels: *in vitro* characterization and *in vivo* assessment for cartilage tissue engineering, J. R. Soc. Interface. 8 (2011) 540–554. 10.1098/rsif.2010.0455.

[30] K.R. Hixon, C.T. Eberlin, P.U. Kadakia, S.H. McBride-Gagyi, E. Jain, S.A. Sell, A comparison of cryogel scaffolds to identify an appropriate structure for promoting bone regeneration, Biomed. Phys. Eng. Express 2 (2016) 035014. 10.1088/2057-1976/2/3/035014.

[31] P. Salmon, D. Trono, Production and Titration of Lentiviral Vectors, CP Human Genetics 54 (2007). 10.1002/0471142905.hg1210s54.

[32] R.A. Heyman, E. Borrelli, J. Lesley, D. Anderson, D.D. Richman, S.M. Baird, R. Hyman, R.M. Evans, Thymidine kinase obliteration: creation of transgenic mice with controlled immune deficiency., Proc. Natl. Acad. Sci. U.S.A. 86 (1989) 2698–2702. 10.1073/pnas.86.8.2698.

[33] M.J. Gardner, S.M. Putnam, A. Wong, P.N. Streubel, A. Kotiya, M.J. Silva, Differential fracture healing resulting from fixation stiffness variability: a mouse model, Journal of Orthopaedic Science 16 (2011) 298–303. 10.1007/s00776-011-0051-5.

[34] M. Orth, M. Braun, C. Scheuer, J. Holstein, A. Khalil, X. Yu, W. Murphy, T. Pohlemann, M. Laschke, M. Menger, BMP-2-coated mineral coated microparticles improve bone repair in atrophic non-unions, eCM 33 (2017) 1–12. 10.22203/eCM.v033a01.

[35] V.J. Alentado, A.M. Knox, C.A. Staut, A.C. McGuire, J.R. Chitwood, S.L. Mostardo, M.Z. Shaikh, R.J. Blosser, U.C. Dadwal, T.-M.G. Chu, C.D. Collier, J. Li, Z. Liu, M.A. Kacena, R.M. Natoli, Validation of the modified radiographic union score for tibia fractures (mRUST) in murine femoral fractures, Front. Endocrinol. 13 (2022) 911058. 10.3389/fendo.2022.911058.

[36] C.B. Erickson, R. Hill, D. Pascablo, G. Kazakia, K. Hansen, C. Bahney, A timeseries analysis of the fracture callus extracellular matrix proteome during bone fracture healing, JoLS (2022) 1–30. 10.36069/JoLS/20220601.

[37] D. Malhan, K. Schmidt-Bleek, G.N. Duda, T. El Khassawna, Landscape of Well-Coordinated Fracture Healing in a Mouse Model Using Molecular and Cellular Analysis, IJMS 24 (2023) 3569. 10.3390/ijms24043569.

[38] W. Huang, C. Zhou, Y. Yu, S. Qin, L. Chen, H. Lin, S. Zhang, L. Xia, W. Liang, Functionalized mesenchymal stem cells for enhanced bone regeneration: advances and challenges, Stem Cell Res Ther 16 (2025) 600. 10.1186/s13287-025-04699-3.

[39] M. Manassero, J. Paquet, M. Deschepper, V. Viateau, J. Retortillo, M. Bensidhoum, D. Logeart-Avramoglou, H. Petite, Comparison of Survival and Osteogenic Ability of Human Mesenchymal Stem Cells in Orthotopic and Ectopic Sites in Mice, Tissue Engineering Part A 22 (2016) 534–544. 10.1089/ten.tea.2015.0346.

[40] M.A. Scott, B. Levi, A. Askarinam, A. Nguyen, T. Rackohn, K. Ting, C. Soo, A.W. James, Brief Review of Models of Ectopic Bone Formation, Stem Cells and Development 21 (2012) 655–667. 10.1089/scd.2011.0517.

[41] L. Mao, D. Zhang, Z. Shen, X. Wang, C. Lai, F. Chen, C. Liu, Recombinant human bone morphogenetic protein-2–engineered piezoplatform synergistically promotes bone regeneration through bone morphogenetic protein receptor activation, Biomaterials Translational 6 (2025) 450. 10.12336/bmt.25.00019.

[42] C.G. Wilson, F.M. Martín-Saavedra, N. Vilaboa, R.T. Franceschi, Advanced BMP Gene Therapies for Temporal and Spatial Control of Bone Regeneration, J Dent Res 92 (2013) 409–417. 10.1177/0022034513483771.

[43] S.N. Lissenberg-Thunnissen, D.J.J. De Gorter, C.F.M. Sier, I.B. Schipper, Use and efficacy of bone morphogenetic proteins in fracture healing, International Orthopaedics (SICOT) 35 (2011) 1271–1280. 10.1007/s00264-011-1301-z.

[44] T.-J. Cho, L.C. Gerstenfeld, T.A. Einhorn, Differential Temporal Expression of Members of the Transforming Growth Factor β Superfamily During Murine Fracture Healing, Journal of Bone and Mineral Research 17 (2002) 513–520. 10.1359/jbmr.2002.17.3.513.

[45] F.J. O’Brien, B.A. Harley, I.V. Yannas, L. Gibson, Influence of freezing rate on pore structure in freeze-dried collagen-GAG scaffolds, Biomaterials 25 (2004) 1077–1086. 10.1016/S0142-9612(03)00630-6.

[46] C.M. Murphy, M.G. Haugh, F.J. O’Brien, The effect of mean pore size on cell attachment, proliferation and migration in collagen–glycosaminoglycan scaffolds for bone tissue engineering, Biomaterials 31 (2010) 461–466. 10.1016/j.biomaterials.2009.09.063.

[47] Z. Jiang, Z. Wu, D. Deng, J. Li, X. Qi, M. Song, Y. Liu, Q. Wu, X. Xie, Z. Chen, Z. Tang, Improved Cytocompatibility and Reduced Calcification of Glutaraldehyde-Crosslinked Bovine Pericardium by Modification With Glutathione, Front. Bioeng. Biotechnol. 10 (2022) 844010. 10.3389/fbioe.2022.844010.

[48] D. Wang, H. Jiang, J. Li, J. Zhou, S. Hu, Mitigated calcification of glutaraldehyde-fixed bovine pericardium by tannic acid in rats, Chinese Medical Journal 121 (2008) 1675–1679. 10.1097/00029330-200809010-00017.

[49] X. Li, H.-Y. Yang, C.M. Giachelli, BMP-2 promotes phosphate uptake, phenotypic modulation, and calcification of human vascular smooth muscle cells, Atherosclerosis 199 (2008) 271–277. 10.1016/j.atherosclerosis.2007.11.031.

[50] C. Lee, H.-G. Lim, C.-H. Lee, Y.J. Kim, Effects of glutaraldehyde concentration and fixation time on material characteristics and calcification of bovine pericardium: implications for the optimal method of fixation of autologous pericardium used for cardiovascular surgery, Interact CardioVasc Thorac Surg (2016) ivw356. 10.1093/icvts/ivw356.

[51] S. Shin, D.M. Salvay, L.D. Shea, Lentivirus delivery by adsorption to tissue engineering scaffolds, J Biomedical Materials Res 93A (2010) 1252–1259. 10.1002/jbm.a.32619.

[52] N. Huebsch, P.R. Arany, A.S. Mao, D. Shvartsman, O.A. Ali, S.A. Bencherif, J. Rivera-Feliciano, D.J. Mooney, Harnessing traction-mediated manipulation of the cell/matrix interface to control stem-cell fate, Nature Mater 9 (2010) 518–526. 10.1038/nmat2732.

[53] N.E. Holmes, P.G.P. Charles, Safety and Efficacy Review of Doxycycline, Clinical Medicine. Therapeutics 1 (2009) CMT.S2035. 10.4137/CMT.S2035.

[54] K.D.N. Gomes, A.P.N.N. Alves, P.G.P. Dutra, G.S.D.B. Viana, Doxycycline induces bone repair and changes in Wnt signalling, Int J Oral Sci 9 (2017) 158–166. 10.1038/ijos.2017.28.

[55] K.N. Agwuh, Pharmacokinetics and pharmacodynamics of the tetracyclines including glycylcyclines, Journal of Antimicrobial Chemotherapy 58 (2006) 256–265. 10.1093/jac/dkl224.

