## Supplemental Figures for "Scaffold-Mediated Cellular and Acellular Delivery of Inducible BMP2-Lentivirus Promotes Atrophic Nonunion Fracture Healing without Heterotopic Ossification"

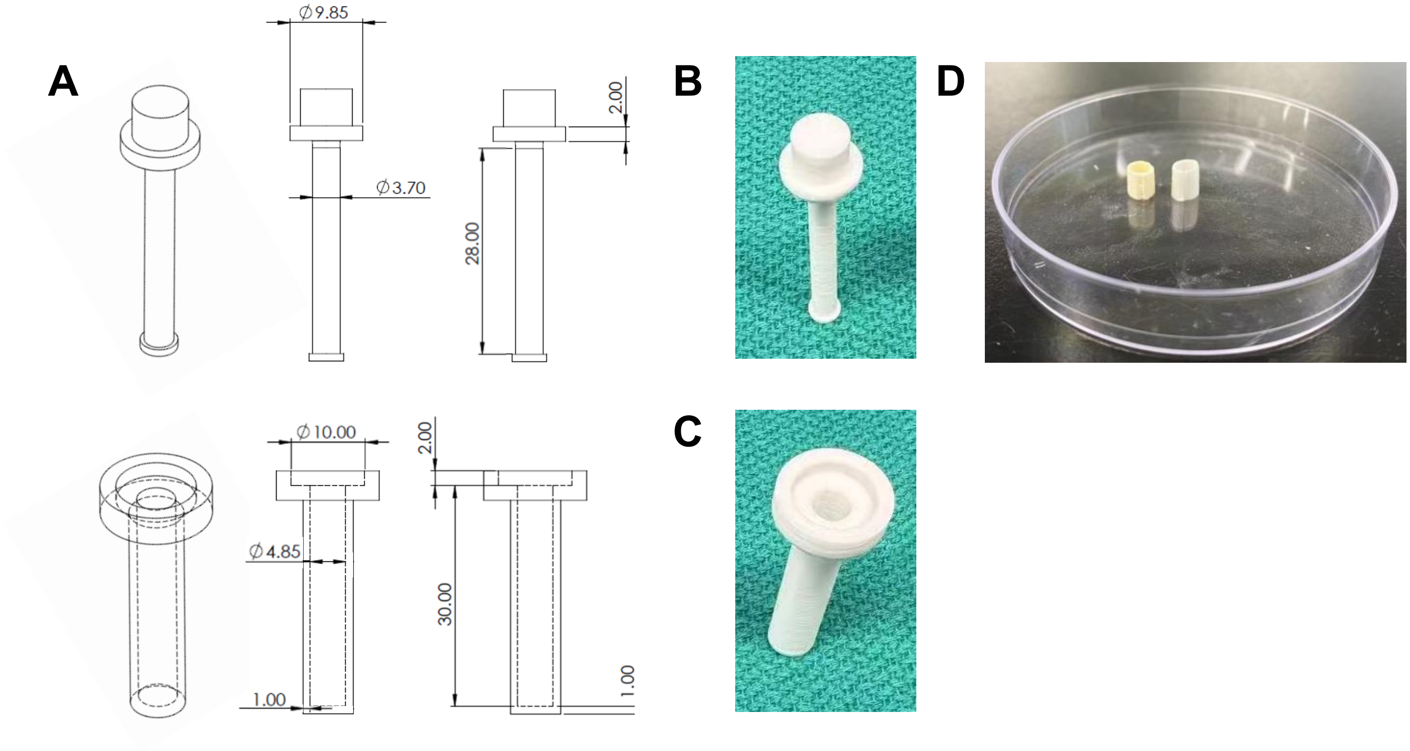


**Figure S1. Custom two-part mold for casting tube-shaped cryogel scaffolds. (A)** Computer-aided design drawings of the two mold components. The molds were made up of polylactic acid (PLA). The inner mandrel (top) consists of a Ø9.85mm flange, 2.00mm thick, and a Ø3.70mm shaft, 28.00mm in length. The outer sleeve (bottom) consists of a Ø10.00mm flange, 2.00mm thick, and a 30.00mm shaft with a Ø4.85mm internal bore and 1.00mm wall thickness. All dimensions are in millimeters. When assembled, the mandrel is seated concentrically within the sleeve. **(B)** 3D-printed inner mandrel and **(C)** outer sleeve. **(D)** Representative CG (left) and CAG (right) cryogel scaffolds cast in the assembled mold following cryogelation at −20°C.

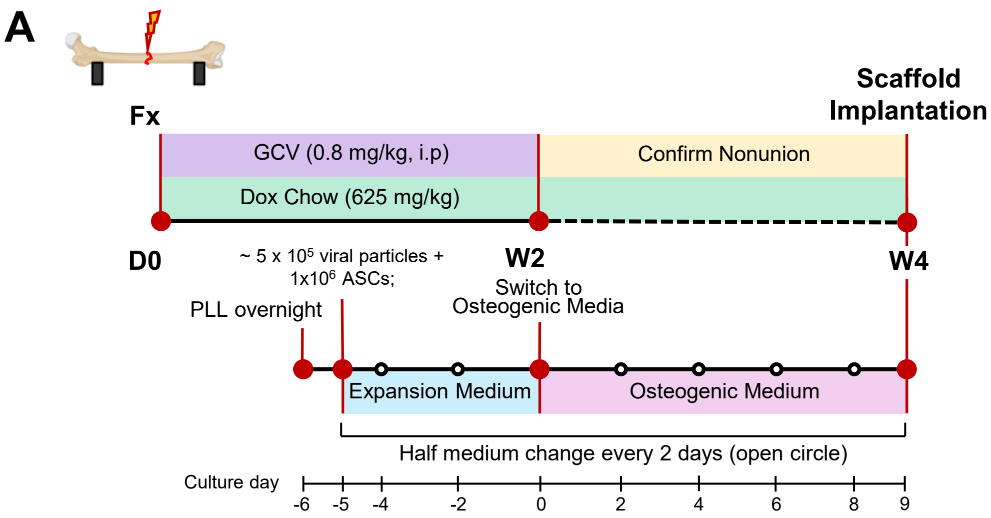


**Figure S2. Experimental timeline of lentiviral transduction of cellular scaffolds.** Scaffolds were coated with poly-L-lysine (PLL) one day before cell seeding. Then, cellular scaffolds were cultured with expansion medium for 5 days before switching to osteogenic medium. All the media was half changed every 2 days.

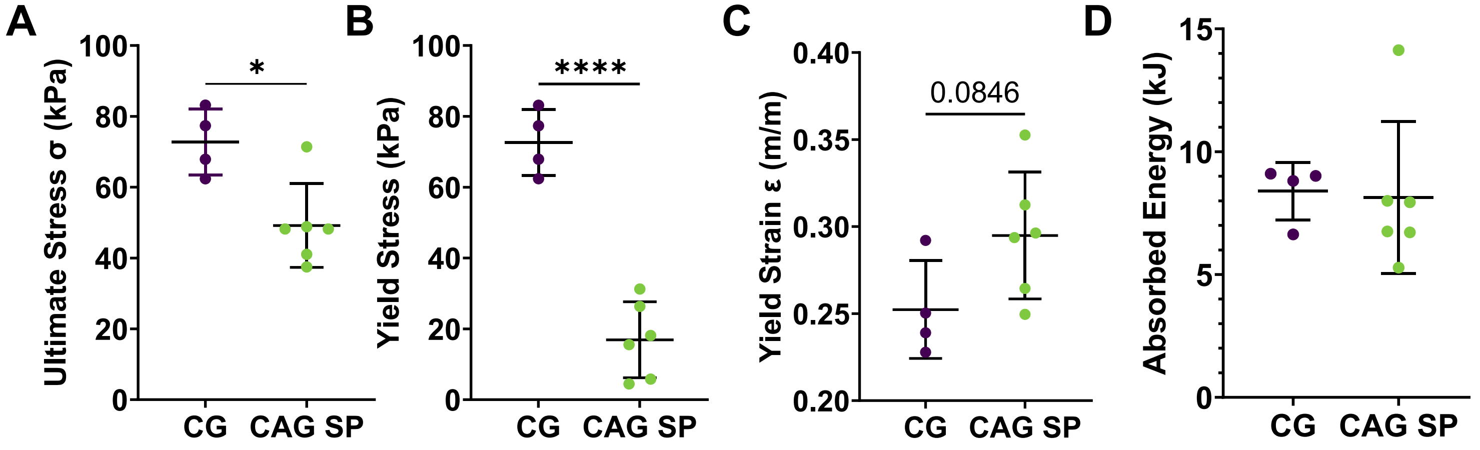


**Figure S3. Mechanical properties of chitosan-gelatin (CG) and chitosan-agarose-gelatin (CAG) cryogel scaffolds. (A)** ultimate stress, **(B)** yield stress, **(C)** yield strain, and **(D)** absorbed energy of CG (n=4) and CAG-SP (n=6) scaffolds. SP denotes small pore formulations. Data are mean ± SD. Statistical comparisons were performed by unpaired t-test *$p < 0.05$, ****$p < 0.0001$.

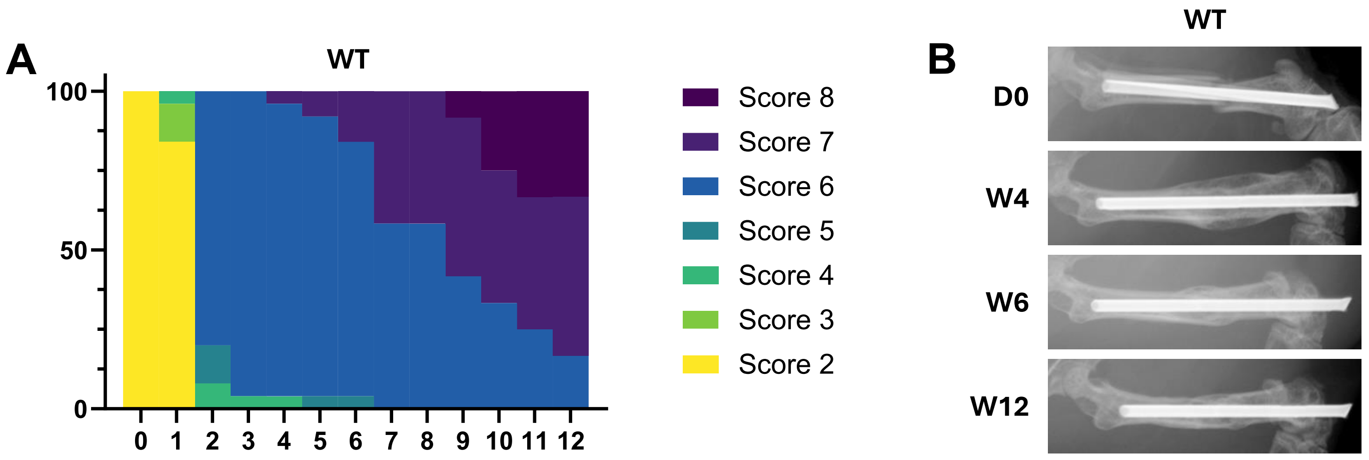


**Figure S4. Femur fracture healing of wild type mice receiving ganciclovir (GCV) was not interfered. (A, B)** Weekly mRUST distributions and representative radiographs at day 0, week 4, 6, and 12 for wild type mice. Stacked bars show the proportion of animals at each score; colors correspond to scores 2-8 as indicated (n =5-10/group).
